# Influenza A M2 Channel Oligomerization is Sensitive to its Chemical Environment

**DOI:** 10.1101/2021.05.07.443160

**Authors:** Julia A. Townsend, Henry M. Sanders, Amber D. Rolland, Chad K. Park, Nancy C. Horton, James S. Prell, Jun Wang, Michael T. Marty

## Abstract

Viroporins are small viral ion channels that play important roles in the viral infection cycle and are proven antiviral drug targets. Matrix protein 2 from influenza A (AM2) is the best characterized viroporin, and the current paradigm is that AM2 forms monodisperse tetramers. Here, we used native mass spectrometry and other techniques to characterize the oligomeric state of both the full-length and transmembrane domain (TM) of AM2 in a variety of different pH and detergent conditions. Unexpectedly, we discovered that AM2 formed a range of different oligomeric complexes that were strongly influenced by the local chemical environment. Native mass spectrometry of AM2 in nanodiscs with different lipids showed that lipids also affected the oligomeric states of AM2. Finally, nanodiscs uniquely enabled measurement of amantadine binding stoichiometries to AM2 in the intact lipid bilayer. These unexpected results reveal that AM2 can form a wider range of oligomeric states than previously thought possible, which may provide new potential mechanisms of influenza pathology and pharmacology.

**Significance Statement:** Many viruses contain small ion channels called viroporins that play diverse roles in viral infections. Influenza A M2 (AM2) is the best characterized viroporin and the target of the antivirals amantadine and rimantadine. Although past structural studies showed AM2 was a monodisperse tetramer, we discovered that AM2 can form polydisperse and dynamic oligomers that are sensitive to their local chemical environment. Our findings provide a new perspective on the structure and mechanisms of AM2 that may extend to other viroporins.

## Introduction

Viroporins are a class of small transmembrane proteins that oligomerize to form channels in membranes.^1^ Found in a range of different viruses, they are involved at multiples stages of infection, including uncoating, replication, assembly, and budding^2,3^ Matrix protein 2 from influenza A (AM2) is a multifunctional viroporin and a clinically approved drug target for amantadine and rimantadine.^3-5^ AM2 is made up of three regions, the extracellular domain, the transmembrane (TM) domain, and the cytosolic tail (Figure 1A). The 20-residue single-pass TM domain of AM2 is necessary and sufficient for oligomerization and formation of a pH-mediated ion channel.^3,6,7^ There are several dozen X-ray or NMR structures of the AM2 TM domain in a variety of membrane mimetics, all depicting monodisperse homotetramers.^8-11^ Despite the uniform oligomeric state, there are significant differences among many of the AM2 structures, and the membrane mimetic used to solubilize AM2 can have major influences on its structure.^9,12^ However, traditional structural biology techniques are limited in their ability to study oligomeric polydispersity, so these existing structures may not capture the full range of possible states. Indeed, earlier fluorescence resonance energy transfer studies suggested that the dimer might be the minimal proton-conducting unit for the full-length AM2 in cells.^13^

**Figure 1.**
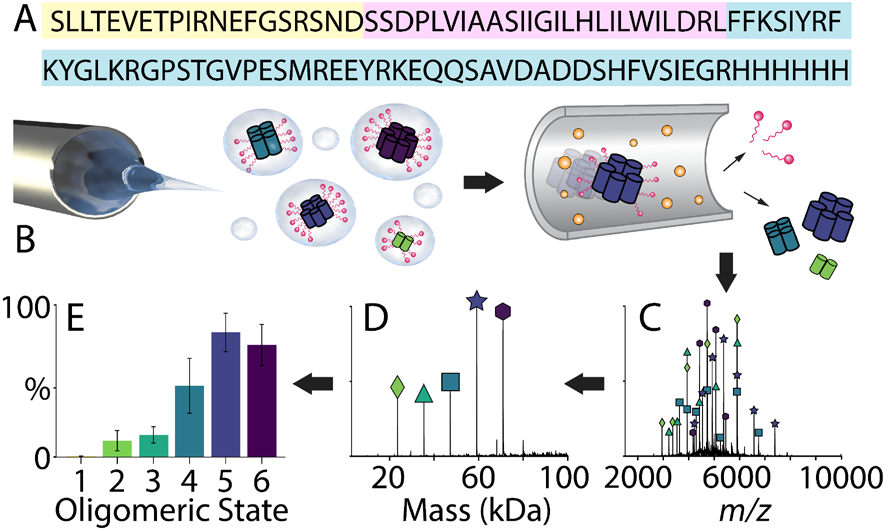
Native MS reveals the oligomeric state distribution of AM2. (A) The sequence of AM2 with the short extra-viral domain colored in *yellow*, the transmembrane domain in *pink*, and the intra-viral region in *blue*. (B) A schematic of ESI with CID to remove detergent from AM2, (C) the mass spectrum of AM2 (at 50 µM per monomer) in C8E4 detergent at pH 5, (D) the deconvolved mass spectrum, and (E) the extracted normalized peak areas of each oligomeric state.

Native mass spectrometry (MS) has emerged as a powerful technique for studying the oligomerization of membrane proteins.^14-16^ For conventional native MS of membrane proteins, the entire protein-micelle complex is ionized with electrospray ionization (ESI).^14^ The detergent adducts are then removed from the protein using collision induced dissociation (CID), and the mass of the bare membrane protein complex reveals the protein stoichiometry and noncovalent ligands that remain bound (Figure 1). Other membrane mimetics, such as nanodiscs, allow membrane proteins to be solubilized in lipid bilayers during native MS.^14,17,18^ Thus, native MS provides rich information and can capture the polydispersity of membrane proteins in different lipid and detergent environments.

Here, we performed native MS on both the full-length and TM AM2 in detergents and nanodiscs. Based upon the existing structures, we predicted that AM2 would form robust tetramers. However, we discovered that AM2 assembled into a range of oligomeric states from dimer to hexamer. Further investigation showed that the oligomeric state of AM2 was influenced by the membrane environment, solution pH, and drug binding. Together, these results reveal that AM2 could be more polydisperse than previously suggested and more sensitive to its chemical environment.

## Results

### AM2 Oligomerization is Sensitive to Detergent and pH

Our initial goal was to investigate drug binding to AM2 using native MS. Based on prior studies,^3,5,9^ we expected to find a monodisperse AM2 tetramer. However, initial results immediately revealed a more complex oligomeric state distribution. To identify conditions that would promote the formation of a monodisperse tetramer, we performed native MS on full-length AM2 to quantify the oligomeric state distribution (Figure 1) in a range of different conditions. We screened different detergents by exchanging AM2 into solution containing tetraethylene glycol monooctyl ether (C8E4), lauryldimethylamine oxide (LDAO), *n*-octyl-β-D-glucopyranoside (OG), *n*-dodecyl-phosphocholine (DPC), *n*-dodecyl-β-maltoside (DDM), and lauryl maltose neopentyl glycol (LMNG). We selected detergents that have been previously used for AM2 structural biology studies, including OG and DPC,^19-23^ as well as detergents that are commonly-used for native MS, such as C8E4, LDAO, and DDM.^24,25^ LMNG was selected for additional structural diversity. For each detergent, we tested pH 5, 7, and 9, which encompass the pH conditions that have been previously investigated with AM2.^26,27^ Selected spectra are shown in Figure 2 with oligomeric state distributions for all plotted in Figure S1.

**Figure 2.**
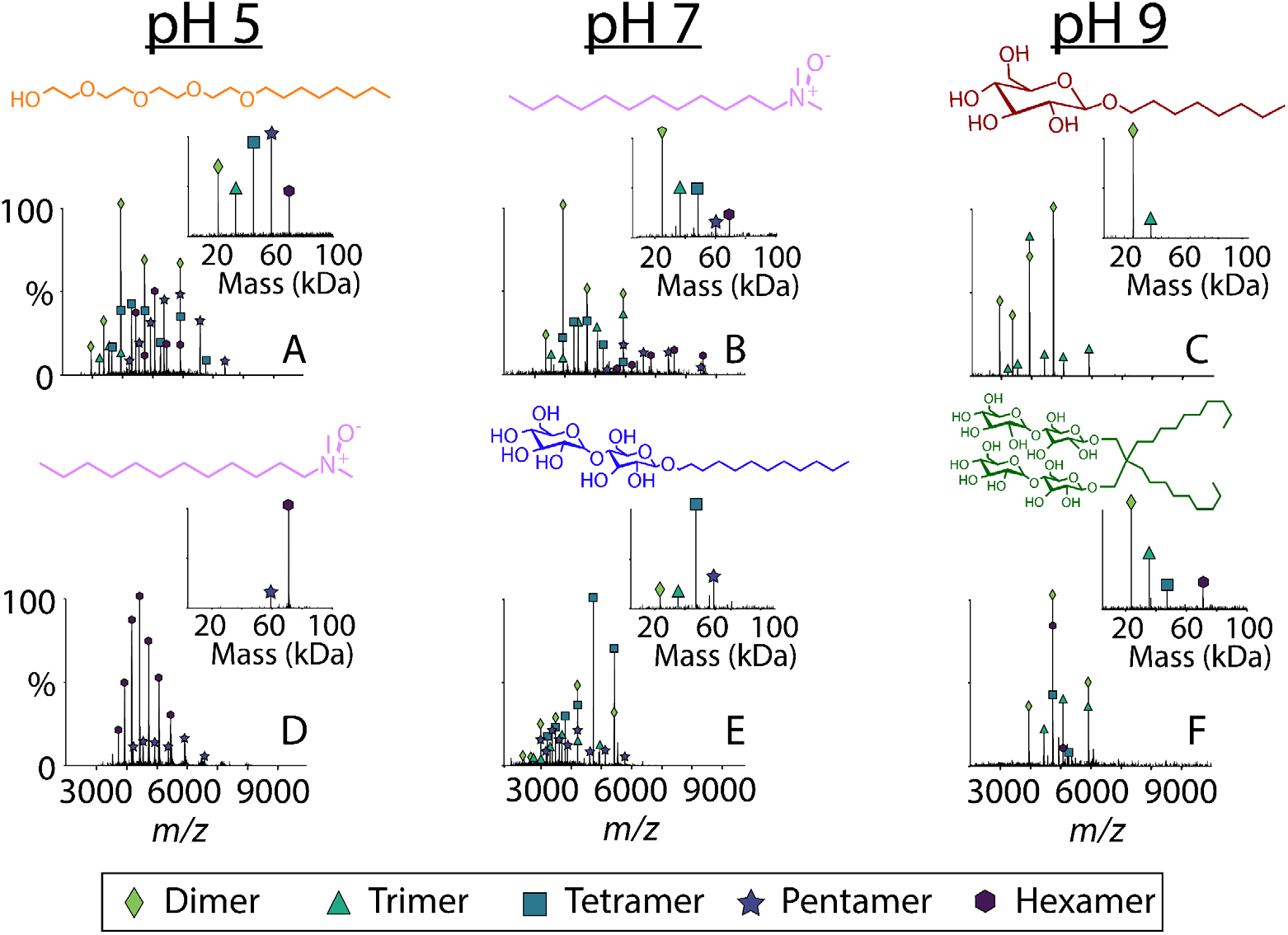
Representative native mass spectra with the deconvolved mass spectra in the inset of AM2 (at 50 µM per monomer) solubilized in (A) C8E4 at pH 5, (B) LDAO at pH 7, (C) OG at pH 9, (D) LDAO at pH 5, (E) DDM at pH 7, and (F) LMNG at pH 9. Each detergent is shown above the spectrum. Average oligomeric state distributions collected in triplicate are shown in Figure S1.

We began by investigating C8E4, which is commonly used for native MS because it is easy to dissociate from membrane proteins.^28,29^ At all pH conditions tested for C8E4, AM2 showed a polydisperse mixture of oligomers that ranged from dimers to hexamers (Figures 2A and S1, Table S1). The precise oligomeric state distribution varied somewhat between replicate measurements, potentially indicating more dynamic oligomers (see Figure S2). Our interpretation is that these more variable oligomers are more sensitive to minor fluctuations in the chemical environment between samples, but the overall trend of forming polydisperse oligomers is highly reproducible. When it was diluted at pH 5, AM2 shifted to lower oligomeric states, indicating weaker interactions in this condition, but it retained higher order oligomers upon dilutions at pH 9 (Figure S3). Overall, AM2 in C8E4 was relatively polydisperse and not heavily influenced by the pH.

In contrast, the oligomeric state of AM2 was more monodisperse and highly dependent on pH when it was solubilized in LDAO. At pH 6 and below, AM2 in LDAO was almost exclusively hexameric, with a small amount of pentamer present (Figures 3A and S4). Additionally, there was almost no variation among replicates of AM2 under this condition, indicating the formation of specific hexameric complexes. However, at pH 7, AM2 in LDAO formed a polydisperse mixture from dimers to hexamers (Figures 2 and 3C). At pH 8 and 9, AM2 was less polydisperse than at neutral pH, forming primarily tetramer with a significant amount of trimer (Figure 3D and E). In contrast with C8E4, these more monodisperse oligomers at pH 5 and 9 remained intact upon dilution, further confirming their specificity (Figure S5). Overall, AM2 formed more selective complexes in LDAO detergent, and the oligomeric states were strongly influenced by the pH.

**Figure 3.**
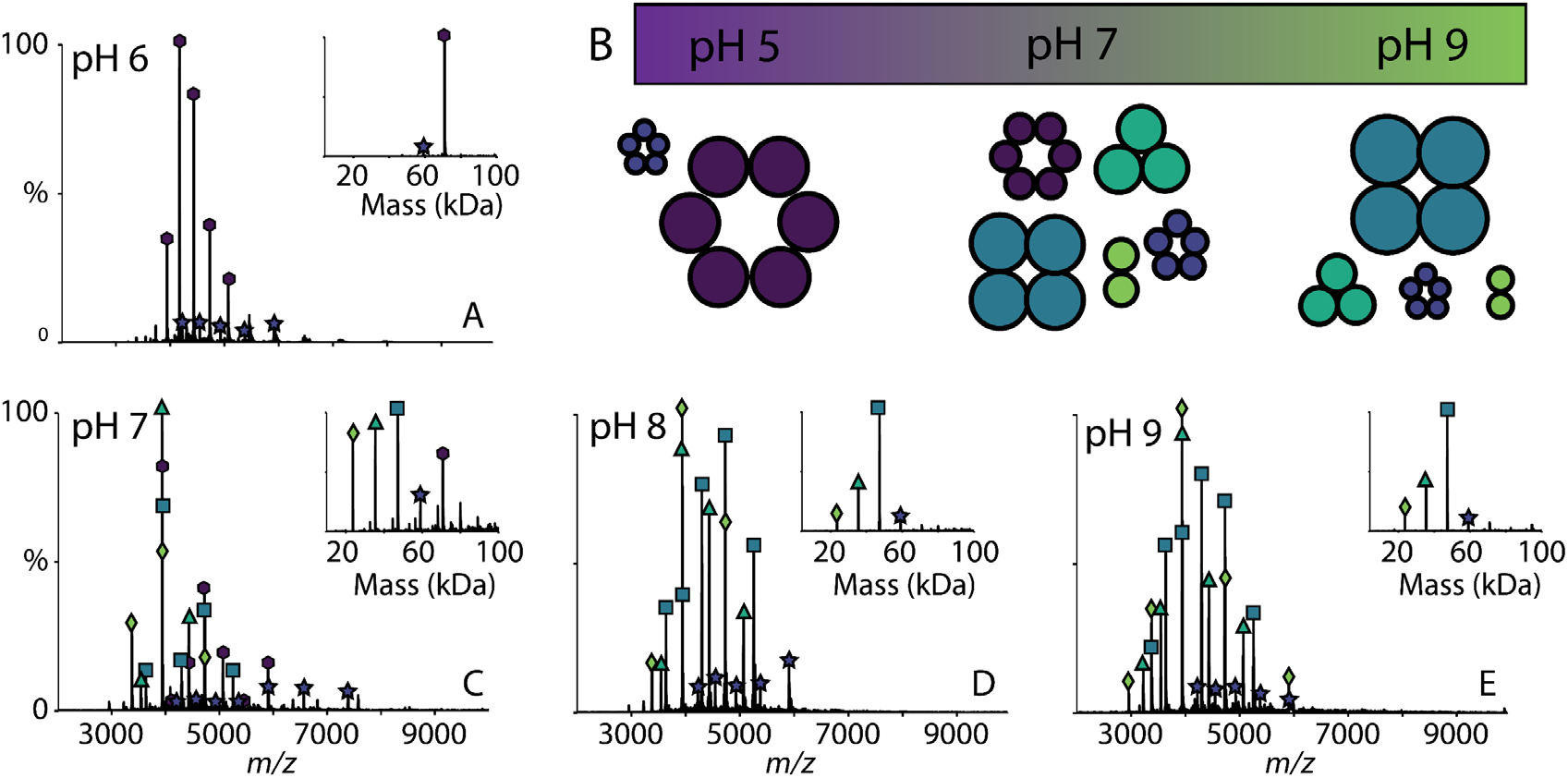
Representative native mass spectra with deconvolved mass spectra (*inset*) of AM2 (at 50 µM per monomer) solubilized in LDAO detergent at pH (A) 6, (C) 7, (D) 8, (E) 9, with (B) a schematic of the different oligomers of AM2 versus pH where the sizes of the oligomers indicate their relative intensities in the spectra. The average oligomeric state distributions collected in triplicate are shown in Figure S3.

The pH also had strong influences on the oligomerization of AM2 in DDM (Figure 4 and S6). At pH 7 and below, AM2 was primarily a mixture of tetramers and pentamers. At pH 9, AM2 in DDM was predominantly trimer with significant amounts of dimer and tetramer (Figure S4). These oligomers also remained intact upon dilution (Figure S6). In contrast, the solution pH did not appear to have a strong influence on the oligomerization of AM2 in OG and DPC detergents (Figure S1). Despite the fact that AM2 has previously been studied in OG and DPC detergents,^21-23,30^ we did not observe monodisperse tetramers, perhaps due to the lower concentrations used here. Instead, there was a general preference for dimer and hexamer. In LMNG, AM2 preferred dimer and trimer at both pH 5 and 9 but was not stable at pH 7 (Figures S1P–Q).

**Figure 4.**
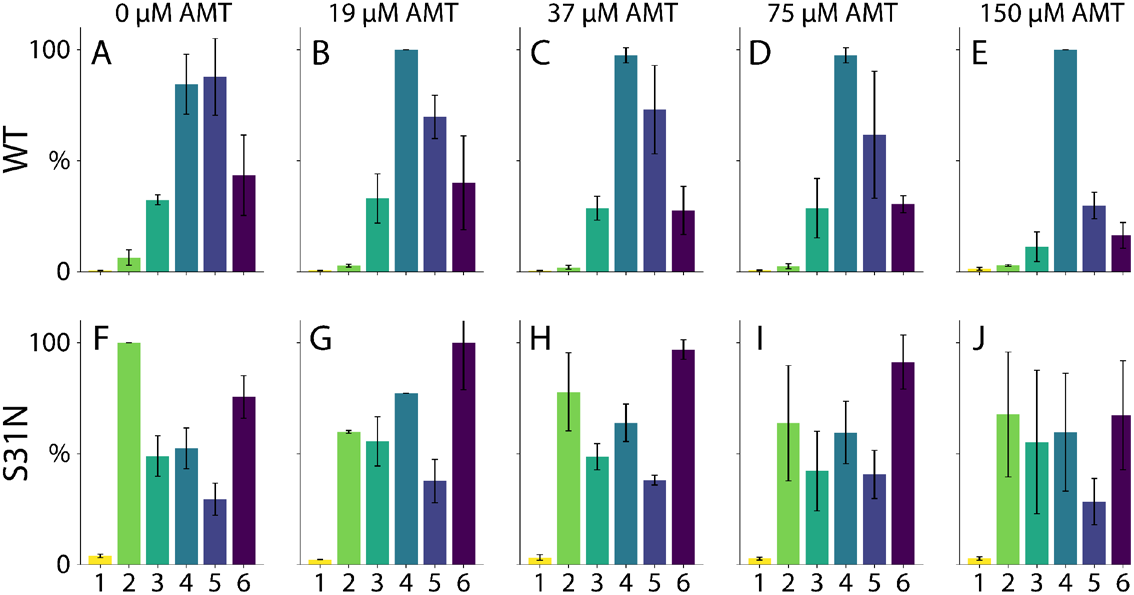
The average oligomeric state of AM2 wild type (A-E) and drug-resistant S31N (both at 50 µM per monomer) (F-J) with 0 µM (A, F), 19 µM (B, G), 37 µM (C, H), 75 µM (D, I), and 150 µM (E, J) amantadine added. Both AM2 WT and S31N were solubilized in C8E4 at pH 9.

Because oligomerization is driven by the transmembrane domain, we next tested the TM domain peptide oligomeric state in select conditions. Similar to the full-length protein, TM-AM2 was polydisperse in C8E4 and OG (Figure S7). In LDAO, TM-AM2 was monodisperse and mostly hexameric at low pH but transitioned to polydisperse above pH 7 (Figure S7). Interestingly, TM-AM2 appeared to have slightly higher preferences for tetramer and hexamer than the full-length AM2 in C8E4 and LDAO detergents. However, the TM peptide overall qualitatively agreed with results from the full-length protein.

Overall, although tetramers were preferred in several conditions, there were no conditions where we found exclusively tetramers (Fig S1 and S7). Instead, we discovered that AM2 oligomerization is influenced by both its detergent environment and solution pH. Depending on the conditions, AM2 can form either highly variable and polydisperse oligomers or relatively selective oligomers of different sizes. Interestingly, the most stable and monodisperse oligomer we found was the hexamer in LDAO under acidic conditions (Figure 3A).

### Orthogonal Measurements Support Oligomeric Variability

Native MS gives accurate relative quantitation for similar species across narrow *m/z* ranges, but differences in ionization, transmission, and detector efficiency make quantitation across wide *m/z* ranges difficult.^31^ To help rule out instrumental biases, we repeated select measurements using a mass spectrometer with a different type of detector. Both the Orbitrap and time-of-flight (ToF) detectors gave similar results (Figures S8 and S9), which support our qualitative conclusions and demonstrates that the results are consistent on different types of mass spectrometers.

We also used ion mobility-mass spectrometry to measure the collisional cross section (CCS) of some of the complexes (Figure S8 and S10).^32,33^ We modeled potential structures assuming oligomerization of the transmembrane domain and disordered soluble domains.^34^ Our experimental CCS values agreed with modeled gas-phase structures, where the disordered regions collapse. Our results also matched predicted CCS values for globular proteins of a similar size,^33^ and the observed charge states are also consistent with a compact structure. Together, these results point to compact oligomers consistent with oligomerization in the transmembrane domain. Based on the observed charge states and CCS values, we can rule out highly extended oligomeric structures and also rule out gas-phase dissociation, which would cause unfolding of the complex and higher CCS values. Also, we would expect any dissociation or complex disruption during native MS to yield a significant population of monomers, which are generally absent. Thus, there is no evidence for complexes being disrupted during native MS. In our interpretation of the data, we have been careful to avoid any conclusions that could be distorted by different ionization efficiencies.

Although both instruments showed similar oligomeric state distributions, we cannot rule out differences in ionization efficiency that could skew the distribution measured by native MS. To further confirm our results, we performed size-exclusion chromatography (SEC) with AM2 in select conditions. It is challenging to directly compare the elution times between different detergents because the micelle sizes can vary. However, qualitative comparisons of the chromatograms of AM2 in different conditions supported the native MS results. Conditions with a wide range of oligomers in native MS, such as C8E4 at pH 9, had broader SEC peaks and more variability between replicate injections (Figure S11). In conditions where AM2 was more monodisperse, such as LDAO at pH 5, we saw narrower and more reproducible peaks.

Similarly, analytical ultra-centrifugation (AUC) was also performed on full-length AM2 in LDAO and C8E4 at pH 5. AUC trends were consistent with native MS. The more polydisperse sample (C8E4) showed several species with AUC, while the more monodisperse sample (LDAO) showed one single species (Figure S12). Thus, these data help support the qualitative descriptions of the oligomeric state distributions and also show changes in the size and polydispersity of the complex in response to the chemical environment. Together, these orthogonal measurements support the qualitative conclusions from native MS.

### Drug Binding Can Remodel AM2 Oligomers

We next measured the effects of amantadine, a clinically-approved inhibitor of AM2,^35^ by adding the drug at different concentrations in all the detergent and pH conditions. Interestingly, we discovered a shift in the oligomerization when amantadine was added to AM2 in C8E4 at pH 9. At low concentrations of amantadine, AM2 formed a range of variable oligomers. At higher concentrations of amantadine, AM2 shifted towards relatively monodisperse tetramers (Figure 4). A similar trend was observed on the ToF platform (Figure S13). We also compared the drug-resistant S31N mutant of AM2 under the same conditions.^36^ Even at high concentrations of amantadine, there were no major changes in the oligomeric state of AM2 S31N.

The S31N mutant appeared to have a similar oligomeric state distribution without added drug (Figure S14). Further experiments in a range of different detergents, pH conditions, and with the full-length and TM peptides of the S31N mutant revealed an overall qualitatively similar oligomeric state pattern (Figure S15). The S31 mutant was generally polydisperse in most conditions but formed monodisperse hexamers in LDAO at pH 5. However, there was some bias towards dimer, suggesting the mutation may affect the oligomeric state distribution in some conditions.

One important limitation of these experiments is that we only observed shifts in the oligomeric state distribution in C8E4 detergent at pH 9. It has been previously found that amantadine preferentially binds under basic conditions, so it is not surprising that we only measured changes at higher pH.^37^ The lack of response in other detergents may be because these detergents cause AM2 to form oligomers with lower drug binding affinity or oligomers with stronger protein-protein interactions that are not easily altered by the drug. AM2 shows the least oligomeric specificity in C8E4, so this set of conditions is perhaps most susceptible to shifts in the oligomeric state distribution caused by the drug.

Another limitation is that only very small signals for drug bound to AM2 were observed, despite the high concentrations added and clear shifts in the oligomeric state distribution induced by drug binding. The lack of signal from bound drug is likely due to gas-phase dissociation of the drug inside the mass spectrometer, where the activation required to remove the detergent micelle also likely removes the small (151 Da) bound drug. Thus, we cannot comment on whether drug is binding in detergents, only on changes in observed oligomeric state as drug is added. Previous work by Pielak *et al*. suggested that amantadine may not be able to bind to AM2 under certain detergent conditions, such as in DHPC micelles, so detergents may be affecting drug binding. In any case, many AM2 structures have amantadine or an analogous AM2 inhibitor added, and our data suggest that the addition of inhibitors may help stabilize the monodisperse tetramer.^19,38,39^

### AM2 in Nanodiscs Shows Lipid Sensitivity and Drug Binding

After screening AM2 in a range of detergent and pH conditions, we characterized its oligomerization in lipid bilayers by assembling AM2 into nanodiscs of different lipid types at a 4:1 ratio of AM2 per nanodisc. Using the shifts in the overall mass of the nanodisc measured by native MS, as well as mass defect analysis (Table S2), we determined the stoichiometry of AM2 embedded within the intact nanodiscs.^40,41^ We first incorporated AM2 into 1,2-dimyristoyl-*sn*-glycero-3-phosphocholine (DMPC) nanodiscs, which showed AM2 stoichiometries from two through six (Figure 5A). We then incorporated AM2 into 1,2-dimyristoyl-*sn*-glycero-3-phosphorylglycerol (DMPG) nanodiscs, which showed less incorporation for the AM2 and only stoichiometries of one, two, or three within the nanodisc (Figure 5B). In both lipids, AM2 had a non-selective distribution of oligomers. In contrast, when AM2 was incorporated into 1,2-dipalmitoyl-*sn*-glycero-3-phosphocholine (DPPC) nanodiscs, it incorporated with stoichiometries of only one and four, which shows that AM2 forms specific tetramers in DPPC bilayers under these conditions (Figure 5D). The increased oligomeric specificity in DPPC nanodiscs may be due to the increased thickness or saturation of the lipid bilayer.^42,43^

**Figure 5.**
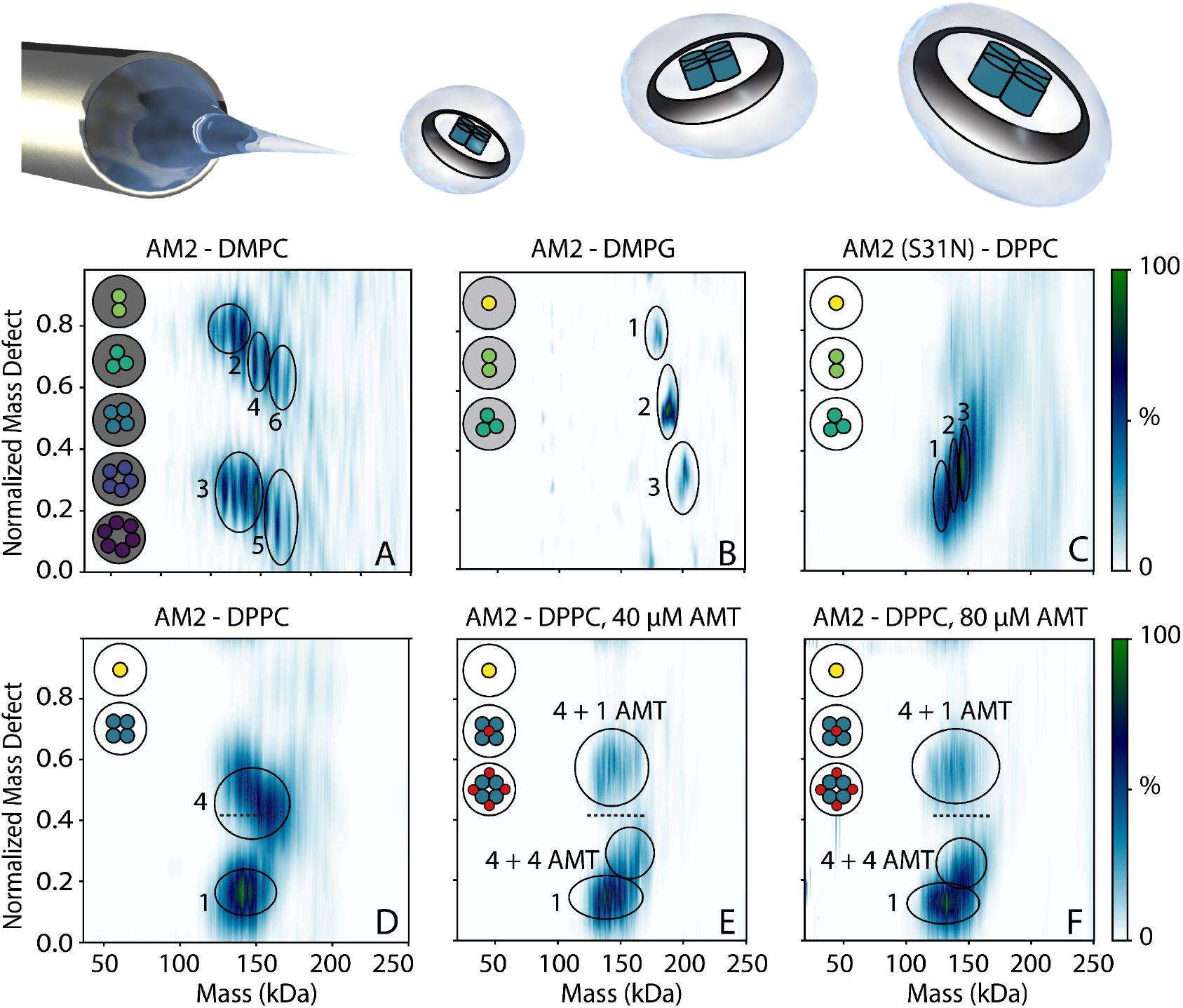
Native MS intensities as a function of normalized mass defect versus mass for (all except C) wild type and (C) S31N AM2 in nanodiscs with (A) DMPC, (B) DMPG, (C–F) DPPC lipids. (E) 40 µM and 80 µM amantadine (AMT) were added, and shifts of the tetramer from the dashed reference line indicate 1 or 4 AMT bound. Illustrations to the upper left indicate observed stoichiometries, which are circled and annotated. The cartoon shown above shows a schematic of directly ionizing intact AM2 nanodiscs.

**Figure 6:**
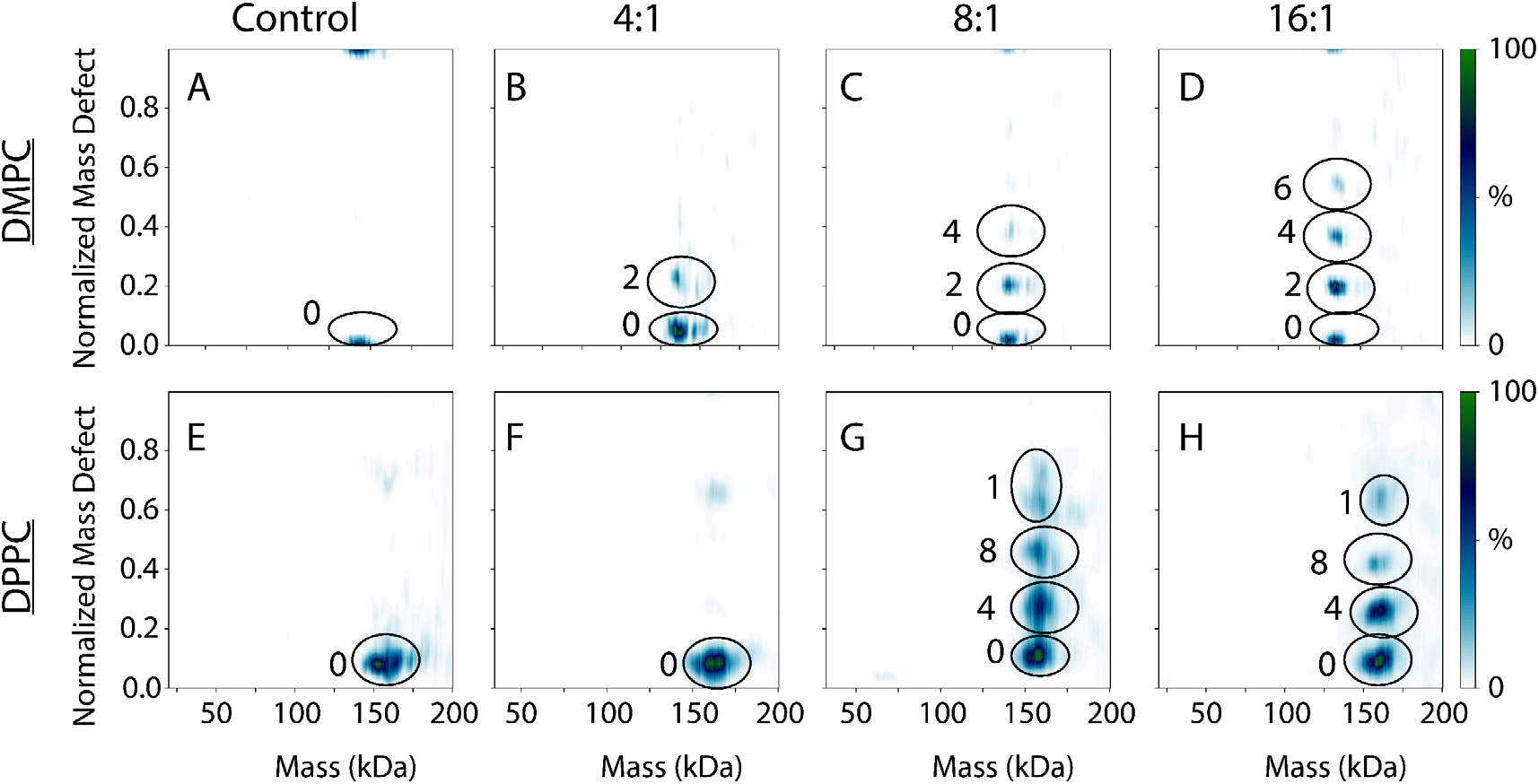
Native MS intensities as a function of normalized mass defect versus mass for the WT TM-AM2 in DMPC nanodiscs (A–D) and DPPC nanodiscs (E–H), with no TM-AM2 added (A, E), a 4:1 ratio (B, F), an 8:1 ratio (C, G), and a 16:1 ratio of TM-AM2 to nanodisc.

We next added amantadine to the DPPC nanodiscs and measured drug binding by native MS. Without amantadine, there were two clear mass defect distributions for monomer and tetramer, respectively. Upon adding 40 µM amantadine, the mass defect of the monomer did not shift, confirming that monomeric AM2 did not bind the drug. However, there were clear shifts in the mass defect of nanodiscs with AM2 tetramers. The first shift corresponded to AM2 tetramers with one amantadine bound (Figure 5E and Table S3). Interestingly, there was also a second shift in the mass defect that corresponded to AM2 tetramer with four amantadine bound. At 80 µM amantadine, the relative intensity of the single-bound state diminished, and the four-bound state became more abundant. DMPC nanodiscs also showed shifts characteristic of drug binding, but the more complex oligomeric state distribution prevented conclusive assignments.

These data agree with existing structures that show AM2 can have one drug bound at lower concentration and four drugs bound at higher concentrations.^39,44^ Specifically, the allosteric binding site located at the helix interface has been previously shown by solution NMR.^39^ Surface plasmon resonance experiments further demonstrated the coexistence of pore binding and allosteric binding sites in AM2.^44^ Recent high-resolution X-ray crystal structures showed that amantadine binds specifically to the pore of the AM2 channel at a one drug per channel ratio at low drug concentrations.^21^ Additionally, at high drug concentrations, rimantadine, an amantadine analog, also binds non-specifically to the AM2 helix interface at a four drug per channel ratio.^44^ Overall, our results from native MS are consistent with prior literature describing binding of amantadine to AM2 in first a 1:4 and a 4:4 ratio, with the later more prevalent at high concentration.

To confirm specificity of drug binding, we incorporated drug-resistant AM2 S31N into DPPC nanodiscs (Figure 5C). AM2 S31N assembled into DPPC nanodiscs in stoichiometries of one, two, and three, suggesting that the mutant did not form specific complexes. Thus, the oligomerization of AM2 S31N appears to be different from the wild type in nanodiscs (Figures 4 and 5). Importantly, AM2 S31N nanodiscs did not show any mass defect shifts upon addition of amantadine, confirming specificity of drug binding (Figure S16).

### AM2 TM Domain Behavior in Nanodiscs

Finally, we investigated the oligomerization of TM-AM2 in lipid nanodiscs by directly adding TM-AM2 to pre-formed nanodiscs. With increasing concentrations of TM-AM2 in DMPC nanodiscs, we measured a mixture of zero, two, four, and six TM-AM2 incorporated into the nanodisc. There have been previous studies of TM-AM2 where has been observed as a dimer of dimers,^45^ so it is not surprising that TM-AM2 incorporated in units of two in the nanodisc. Our TM-AM2 results also differed from the more random pattern of incorporation that we measured with the full-length AM2. The difference between the full-length and TM AM2 reveals that the disordered cytosolic region of the full-length AM2 may influence the oligomerization of AM2 within DMPC lipid bilayers. In contrast, with DPPC nanodiscs, we saw a very similar trend to the full-length AM2, with TM-AM2 being incorporated in units of four but with a small amount of monomer present.

## Discussion

Here, we used native MS to study the oligomerization of full-length and TM-AM2 in different pH conditions, detergents, lipid bilayers, and with added drug. In nearly all the detergent and pH combinations screened, AM2 had different patterns of oligomerization, which reveals two key conclusions. First, AM2 is not exclusively a tetramer. Second, AM2 can be sensitive to its chemical environment, showing different oligomeric states in different pH and lipid/detergent conditions.

There are two potential interpretations of these surprising results. On one hand, it may be that the tetramer is the true physiological state of AM2. In this case, our results reveal that it can be challenging to capture the pure tetramer in detergent and even some lipid bilayers. Native MS thus reveals conditions that favor or disfavor the true physiological oligomer. For example, AM2 has a strong propensity to form tetramers in DPPC nanodiscs. In contrast, our results with OG and DPC detergents do not show monodisperse tetramer as would be expected from past NMR and AUC studies in these detergents.^19-21,23,46^ It could be that differences in protein or detergent concentrations, peptide length, or other experimental conditions caused these discrepancies. Our results generally show more robust tetramer in bilayers over detergents, with drug added, and with the TM peptide over the full-length, so these conditions may favor tetramer. Past research has shown significant changes in structure depending on bilayer/detergent conditions, and these structural changes could go beyond conformation to include changes to the oligomeric state.^45^

However, another interpretation of our results is that the oligomeric states of AM2 are more complex than previously thought. It is very challenging to measure the oligomeric state distribution for small membrane proteins like this, especially if they form polydisperse oligomers.^47^ Past studies may have underestimated the true polydispersity due to limitations of the analysis techniques. For example, crystallization could push AM2 to form tetramer complexes or select for conditions where structurally monodisperse tetramers are present. Most X-ray structures of AM2 were collected in LCP, which could favor tetramers.^48-50^ It is challenging to directly measure the oligomeric state distribution for homo-oligomers with NMR without advanced techniques that are not always employed.^51^ Furthermore, many structural studies have been conducted in the presence of high drug concentrations, which may bias the drug towards a monodisperse tetramer, as we saw here (Figure 4). Native MS, despite the potential biases outlined above, provides a direct analysis of the oligomeric state distribution of AM2 that could reveal previously unseen oligomers. Past native MS studies have shown similar oligomeric pore-forming proteins, such as the mechanosensitive channel of large conductance (MscL),^52^ also form polydisperse oligomeric complexes that are sensitive to the local chemical environment. Conversely, other native MS studies have shown similarly small oligomeric membrane protein complexes to form specific monodisperse oligomers.^53^

These results could present several new hypotheses for AM2 structure and function in a physiological context. First, AM2 is known to be activated by lower pH.^54^ Our results in LDAO detergent may suggest that this could be aided by shifts in oligomeric state distribution (Figure 3). Other detergents do not show as clear of a shift, but higher oligomers are preferred at lower pH in several different conditions. It may be that AM2 forms smaller oligomers at neutral pH, but acidic conditions in the endosome trigger formation of larger oligomeric pores that cause the influenza virus to fuse with the endosomal membrane and release the nucleic acid cargo for replication.^55^

Our results also suggest that changes in the lipid environment may affect the oligomerization of AM2 (Figure 5). DPPC nanodiscs showed specific tetramers whereas DMPC nanodiscs showed less selective complexes. The thickness and fluidity of the lipid bilayer are known to influence AM2 activity, and these functional changes may be due, in part, to changes in the oligomeric state distribution.^42,56^ Different lipid compositions in different intracellular organelle membranes or between different virus strains may contribute to altering AM2 activity.^57^

Finally, our results propose a new potential mechanism of drug activity where the drug may affect oligomerization. It likely still blocks the channel directly or by inducing conformational changes, but it may have the added effect of altering the oligomeric state distribution. Similar effects of AM2 stabilization by drug binding have also been observed in solution and solid-state NMR studies.^19,58,59^ Clearly, extensive future studies will be required to test all these hypotheses, but our results shed new light on AM2 oligomerization and prompt a fresh perspective on its mechanisms that may extend to other viroporins.

These experiments also mark a technical milestone in using native MS to measure drug binding to a membrane protein in an intact lipid bilayer. High-resolution native MS enabled detection of a 151 Da drug bound to a roughly 150 kDa intact nanodisc complex containing a polydisperse mixture of lipids and AM2. We were able to simultaneously determine the stoichiometry of the bound drug as well as which AM2 oligomer it was binding. Importantly, nanodiscs seemed to better preserve the drug bound complex inside the mass spectrometer than detergent micelles, which were unable to capture much of the bound drug. We suspect that the nanodisc better protects the protein-drug complex by preserving the membrane protein in its surrounding lipid bilayer.

In conclusion, we discovered that AM2 is more polydisperse than previously thought and can be influenced by both the pH and the surrounding membrane environment. In some conditions, AM2 assembles into specific complexes, but others create a dynamic mixture of oligomers. Overall, the application of new analytical approaches revealed unexpected biophysical insights into the polydispersity and pharmacology of AM2 that may have implications for the structures and functions of other viroporins.

## Materials and Methods

### Preparation of AM2 in Different Detergents and pH

Full-length AM2 was expressed and purified as previously described, and details are provided in the Supporting Information. Purity was confirmed by SDS-PAGE and native MS, which both showed no detectable contaminants. Protein activity was confirmed with proton flux assays with POPC liposomes (Figure S17). A series of ammonium acetate solutions were first adjusted to pH 4, 5, 6, 7, 8, and 9 with acetic acid or ammonium hydroxide. All detergents were purchased from Anatrace. Each detergent solution was created by adding twice the critical micelle concentration (CMC) of the detergent to the ammonium acetate solution at each pH. AM2 was exchanged into each of these detergent solutions using Bio-Spin 6 columns (Bio-Rad) and diluted to a final concentration of 50 µM (per monomer) prior to analysis in the relevant solution, except where different concentrations are noted. Samples were allowed to briefly equilibrate at room temperature prior to analysis, but no significant changes were observed in the oligomeric state distributions over time or at colder temperatures. For TM-AM2, the peptide was synthesized as previously described^60^ and diluted to 50 µM in each detergent solution. For drug binding experiments, amantadine (Sigma Aldrich) was diluted to 1.5, 0.75, 0.375, and 0.188 mM in water. 0.5 µL of amantadine was added to 4.5 µL of AM2, for a final drug concentration of 150, 75, 37.5, and 18.8 µM. Mixtures were incubated with amantadine for 5− 10 minutes prior to analysis.

### Nanodisc Assembly and Sample Preparation

AM2 nanodiscs were assembled using a 4:1 AM2 to nanodisc ratio. Lower ratios of incorporation showed less AM2 incorporated and higher ratios showed complex spectra that were difficult to resolve and interpret. For nanodiscs containing DMPC and DMPG lipids, the lipids (Avanti Polar Lipids) solubilized in cholate (Sigma Aldrich) were added at an 80:1 ratio of lipid to membrane scaffold protein (MSP). Details on MSP expression and purification are provided in the Supporting Information. For nanodiscs containing DPPC lipids, the lipids were added at a 90:1 ratio of lipid to MSP. All nanodiscs were assembled overnight by adding Amberlite XAD-2 hydrophobic beads (Sigma Aldrich) at the phase transition temperature of the lipid. To isolate nanodiscs containing AM2 from empty nanodiscs, nanodiscs were purified using a HisTrap HP 1 mL column (GE Healthcare). The column was equilibrated with buffer containing 40 mM Tris, 0.3 M NaCl, and 20 mM imidazole at pH 7.4. AM2 nanodiscs were then eluted from the column with buffer containing 40 mM Tris, 0.3 M NaCl, and 400 mM imidazole at pH 7.4. Nanodiscs were then concentrated and purified on a Superose 6 10/300 GL (GE Healthcare) equilibrated with 0.2 M ammonium acetate. For all nanodisc drug binding experiments, 1 µL of 400 or 800 µM drug were added to 9 µL of nanodiscs for final drug concentrations of 40 and 80 µM. These samples were allowed to incubate for ten minutes at room temperature prior to analysis.

Nanodiscs for peptide experiments were assembled at a 90:1 and 80:1 ratio of lipid to MSP for DPPC and DMPC nanodiscs respectively. All nanodiscs were assembled overnight by adding Amberlite XAD-2 hydrophobic beads (Sigma Aldrich) at the phase transition temperature of the lipid. Nanodiscs were then purified on a Superose 6 10/300 GL (GE Healthcare) equilibrated with 0.2 M ammonium acetate. After purification, all nanodiscs were diluted to a final concentration of µM. Nanodiscs were then mixed with peptide at a 16:1, 8:1, 4:1, 2:1, and 1:1 ratio of peptide to nanodisc and allowed to incubate for 30 minutes at room temperature prior to analysis.

### Native Mass Spectrometry

Native MS was performed using a Q-Exactive HF Orbitrap (Thermo Scientific, Bremen) mass spectrometer with ultra-high mass range modifications except where noted as a Synapt XS Q-ToF mass spectrometer (Waters Corporation, Manchester). The native mass spectra were deconvolved and quantified using UniDec, and macromolecular mass defect analysis was used to quantify the stoichiometries of AM2 and amantadine in nanodiscs.^41,61,62^ Full details are provided in the Supporting Information. Prior published results with streptavidin, a similarly sized tetramer, with similar instrument conditions provided a positive control demonstrating the ability of native MS to preserve and detect specific noncovalent complexes of the same size.^61^ Similar experiments on a small transmembrane protein complex, semiSWEET, also demonstrate the ability of native MS to detect specific complexes of small membrane proteins.^63^

## Supporting information

Response to Reviewers

## Acknowledgments

The pMSP1D1 plasmid was a gift from Stephen Sligar (Addgene plasmid #20061). The authors thank Maria Reinhardt-Szyba, Kyle Fort, and Alexander Makarov at Thermo Fisher Scientific for their support on the UHMR Q-Exactive HF instrument. This work was funded by the National Institute of General Medical Sciences and National Institutes of Health (T32 GM008804 to J.A.T. and R35 GM128624 to M.T.M.). A.D.R. is supported by a National Institutes of Health grant (T32 GM007759) and the Doctoral Dissertation Research Fellowship, Peter O’Day Fellowship in Biological Sciences, and Office of the Vice President for Research and Innovation at the University of Oregon. A.D.R. is an ARCS Foundation Scholar supported by the ARCS Oregon Chapter. J.S.P. thanks the National Institute of Allergy and Infectious Diseases (R21AI125804-02) for generous support. C.K.P. and N.H. are supported by the National Science Foundation (NSF MRI Award No. 2018942). J. W. is supported by the National Institute of Allergy and Infectious Diseases at the National Institute of Health (NIH R21 AI144887 and R33 AI119187).

## Data Availability

Native mass spectra, ion mobility data files, SEC chromatograms, and modeled structures have been deposited in Figshare (DOI: 10.6084/m9.figshare.14558079.v1).

