## Supporting Information for "Influenza A M2 Channel Oligomerization is Sensitive to its Chemical Environment"

### Supplemental Methods

#### Protein Expression and Purification

The full-length AM2 with C-terminal polyhistidine tag and cysteines converted to serines was overexpressed in BL21(DE3) pLysS cells. The sequence is shown in Figure 1A without the N-terminal methionine, which is cleaved during expression. Cells were grown at 37 °C in terrific broth media (Thermo Fisher Scientific) to an optical density of 0.8-1.0. Overexpression was induced with isopropyl β-d-1-thiogalactopyranoside at a final concentration of 1 mM for three hours, and cells were then harvested through centrifugation. Cells were resuspended in lysis buffer containing 150 mM NaCl, 50 mM Tris, 40 mM octyl-glucoside (OG), and protease inhibitor. After resuspension, cells were lysed using the LM20 Microfluidizer High Sheer Homogenizer. Lysed cells were then stirred at 4 °C for 1–3 hours to allow for membrane solubilization. The lysate was clarified through centrifugation at 48,380×*g* for 20 minutes. Prior to protein purification, a HisTrap HP 5 mL column (GE Healthcare) was equilibrated with buffer A (150 mM NaCl, 50 mM Tris, 40 mM OG, 20% glycerol, and 20 mM imidazole). The sample was then filtered, loaded to the column, and washed with 10–15 column volumes of buffer A. To remove any nonspecific protein binding, the column was then washed with 5–10 column volumes of 5% buffer B (150 mM NaCl, 50 mM Tris, 4 mM OG, 20% glycerol, and 300 mM imidazole). AM2 was then eluted with 100% buffer B. It was then diluted with buffer A to a final monomer concentration of 580 μM, aliquoted, and flash frozen. The S31N mutant of AM2 was expressed and purified using the same protocol as AM2 wild type.

Membrane scaffold protein, MSP1D1(–) was expressed and purified as previously described.[1,2] Briefly, MSP1D1 was expressed in *E. coli* and purified using immobilized metal affinity chromatography (IMAC). Following cleavage of the polyhistidine tag, MSP1D1(–) was purified by reverse IMAC.

#### Native Mass Spectrometry

Native mass spectrometry was performed as previously described[3,4] using a Q-Exactive HF Orbitrap (Thermo Scientific, Bremen) mass spectrometer with Ultra-High Mass Range Modifications except where stated otherwise. Nano-electrospray ionization in positive ion mode was performed using borosilicate needles pulled using a P-1000 micropipette puller (Sutter Instruments).

Detergent-solubilized AM2 was analyzed with a range of 1,500−15,000 *m/z* at a resolution of 15,000. The trapping gas pressure was set to 5, and the spray voltage ranged from 1.1−1.5 kV. To aid in desolvation and detergent removal, 10−50 V of higher-energy collisional dissociation (HCD) energy and 10–50 V of source fragmentation were applied to each sample, as previously described.[4] The precise collision voltages were adjusted slightly for each sample, and results are shown for the lowest value that gave a well-resolved spectrum. An open vial with 2–5 mL of acetonitrile was placed in the source of the mass spectrometer to allow for vapor charge reduction of all samples, which we found helped stabilize complexes during native MS.[5] Mass spectrometry data was collected as single measurements for three sets of dilutions after the protein was buffer exchanged. Spectra are shown for a single representative replicate, and error bars show the standard deviation of the three replicates.

Nanodiscs were analyzed with a range of 2,000−25,000 *m/z* at a resolution of 15,000. The trapping gas pressure was set to 5 with a spray voltage of 1.1−1.3 kV. For nanodiscs with 1,2-dimyristoyl-*sn*-glycero-3-phosphocholine (DMPC) or 1,2-dimyristoyl-*sn*-glycero-3-phosphorylglycerol (DMPG) lipids, 50−100 V of HCD collisional energy and 10−50 V of source voltage was applied to aid in the desolvation. To aid in the analysis of 1,2-dipalmitoyl-*sn*-glycero-3-phosphocholine (DPPC) nanodiscs, a super charging reagent, propylene carbonate (Arcos Organics), was added prior to ionization at 5% propylene carbonate by volume.[3] For DPPC nanodiscs, 100−200 V of HCD collisional activation was added to remove propylene carbonate. Representative spectra are shown from three replicate nanodisc assemblies.

Ion mobility-mass spectrometry (IM-MS) analysis was performed on a Synapt XS HRMS Q-ToF mass spectrometer (Waters Corporation, Manchester) using a nano-electrospray ionization source with borosilicate glass capillaries prepared as described above. MS conditions were applied to remove detergent adducts without disrupting structure prior to detection with instrument parameters as follows: capillary voltage, 1.5–1.8 kV; sampling cone, 150 V; trap collision energy, 100 V; transfer collision energy, 10 V; trap gas, 10 mL/min; helium cell gas, 120 mL/min; backing gas, 2.85 mbar. The parameters for IM were as follows: IM cell wave height, 40 V; IM cell wave velocity, 1000 m/s; transfer wave height, 4 V; transfer wave velocity, 69 m/s. Arrival time distributions (ATDs) were viewed using DriftScope 2.9 (Waters Corporation). CCS values were calculated as previously described using standards with published values.[2] All reported CCS values were the result of triplicate experiments, and error bars are shown as the standard deviation of the CCS for different charge states.

**Native MS Data Analysis**

The native mass spectra for AM2 solubilized in detergent were deconvolved using UniDec as previously described.[4] The settings for the deconvolution of AM2 in all conditions included a mass range of 1−110 kDa, a charge range of 1−50, and a FWHM of 1 *m/z*. A curved background subtraction of 100, as well as point smooth width of 1 and a beta value of 50 were also applied for all data.[6,7] The native mass spectra for the AM2 nanodiscs were analyzed using UniDec as previously described.[8] The mass range was extended to an upper limit of 250 kDa. For nanodiscs made of DMPG and DMPC lipids, the charge range was set 1−25. For nanodiscs made of DPPC lipids, the charge range was set 1−16. The mass of the lipid was used with mass smoothing set to -1.

To determine the stoichiometry of both full-length AM2 and TM-AM2 in nanodisc samples, we used mass defect analysis.[8,9] Mass defect analysis divides the mass of the sample by a reference mass (the mass of the lipid), and the remainder of the division is then plotted. The plotted remainder is then normalized between 0 and 1.[8] Nanodiscs with the same number of proteins or peptides associated but varying numbers of lipids incorporated will have the same mass defect. This allows for us to sum the mass defect of the protein or peptide across nanodiscs with different numbers of total lipids, yielding the overall mass defect distribution. Mass defect analysis thus reveals the number of AM2 molecules associated with an intact nanodisc inside the mass spectrometer. The addition of each full-length AM2 incorporated into the nanodisc also shifted the overall mass of the complex by about 10 kDa, which further helped in the assignment of the stoichiometry of the protein-nanodisc complexes.

**AM2 Model Structures and Predicted Collisional Cross Sections**

CCS values expected for native, globular proteins were determined using the empirical relationships between mass and CCS for a variety of proteins.[10,11] Model structures of AM2 oligomers were generated in PyMol[12] using PDBs 2N70[13] and 4N8C[14] as templates for the transmembrane and intracellular domains. Any discrepancies in sequence were changed, and missing extracellular domain residues were added manually in PyMol to generate a model AM2 monomer. This monomeric structure was then subjected to a brief (1 ns) relaxation in water using

GROMACS, and the resulting simulated structure was used to construct all model oligomeric complexes with PDB 2KIH[15] as a template for subunit arrangement.

*In vacuo* molecular dynamics simulations of each model AM2 structure were then performed using the GROMOS96 43a2 force field in GROMACS, as previously described.[16] Briefly, the center of the experimental charge state distribution was chosen for each AM2 oligomer, and a low-energy configuration of positive charges was determined using the charge placement algorithm in Collidoscope.[17] This configuration was used to assign charges during topology file generation, and then each model AM2 structure was allowed a brief energy minimization step, followed by a 5 ns *in vacuo* MD production run at 300 K with a modified Berendsen thermostat. CCSs for simulated structures were computed using nitrogen buffer gas and the Trajectory Method in Collidoscope after identifying a low-energy charge configuration for the compacted structures.

**Size Exclusion Chromatography**

Analytical size exclusion chromatography (SEC) was performed using a Superdex 200 Increase 10/300 (GE Healthcare) equilibrated with 1 column volume of each solution, and 100 µL of concentrated AM2 (580 µM) was injected in duplicate.

**Analytical Ultracentrifugation**

Samples for analytical ultracentrifugation (AUC) were prepared similarly to native MS samples by buffer exchanging full-length AM2 into 0.2 M ammonium acetate at pH 5 with twice the critical micelle concentration of either C8E4 or LDAO detergent with a final protein concentration of 2 mg/ml. Experiments were performed using a Beckman Optima Analytical Ultra Centrifuge with a 4 cell An-60 Ti rotor. After equilibrating 4 hours at 25 °C, samples were spun at 40,000 rpm for sedimentation velocity experiments. The absorbance data was collected at 280 nm every 2 minutes overnight (>12 hours or until the baseline was well established by protein depletion). All AUC data was collected in triplicate from separate spins. Data from absorbances at 280 nm were fit with direct boundary modeling and analyzed using the c(s) distribution in Sedfit.[18] Buffer density was estimated at 1.00 g/L, and the viscosity was assumed to be 1.00 cP. Partial specific volume for the protein was assumed to be 0.73 mL/g.

**Liposome Assays**

Proton flux assays were performed on AM2 by assembling AM2 into liposomes made of 1-palmitoyl-2-oleoyl-*sn*-glycero-3-phosphocholine (POPC) lipids. Liposomes were assembled with 10 mg of POPC solubilized in 0.1 M sodium cholate, 20 nmol of full-length AM2 solubilized in OG detergent, and 0.2 nmol of valinomycin. The internal liposome buffer (50 mM phosphate, 50 mM citrate, 122 mM KCl and 122 mM NaCl) was added to a final mixture volume of 500 µL. Amberlite XAD-2 hydrophobic beads (Sigma Aldrich) were added to the mixture, and liposomes were assembled at 4 °C. The liposomes were then extruded, and the assay was performed as previously described.[15] Solution pH measurements were made using a pH microelectrode (InLab) and measurements were made each second. Assays were performed by adding 2 µL of 1 M HCl to the solution being mixed with a stir bar. An increase in the solution pH after acid was added was only observed in liposomes where both AM2 and valinomycin were present. Controls were performed on liposomes without AM2 and without valinomycin, and there was no proton flux measured in these conditions.

### Supplemental Figures

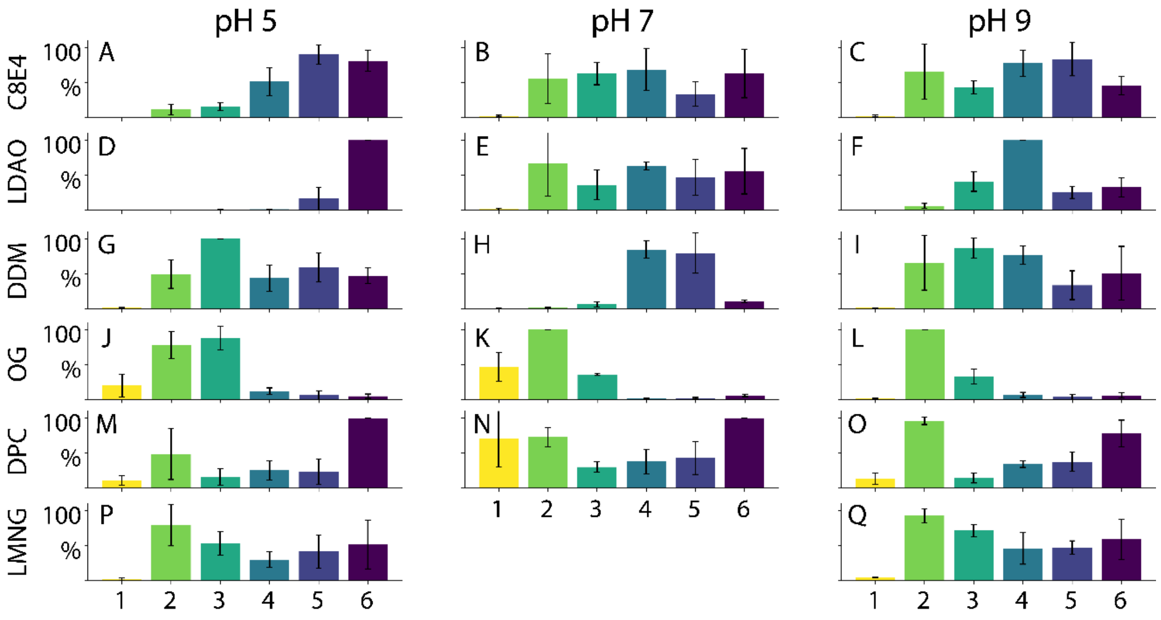

**Fig. S1**: The average relative peak areas measured by native MS of different oligomeric states of WT full-length AM2 (at 50 μM monomer) at pH 5 (A, D, G, J, M, P), pH 7 (B, E, H, K, N), and pH 9 (C, F, I, L, O, Q) while solubilized in C8E4 (A–C), LDAO (D–F), DDM (G–I), OG (J–L), DPC (M–O), and LMNG (P, Q). AM2 was not stable in pH 7 with LMNG and no mass was detected under these conditions. Error bars indicate the standard deviation of measurements from triplicate samples. Representative native mass spectra of select conditions are shown in Figure 2.

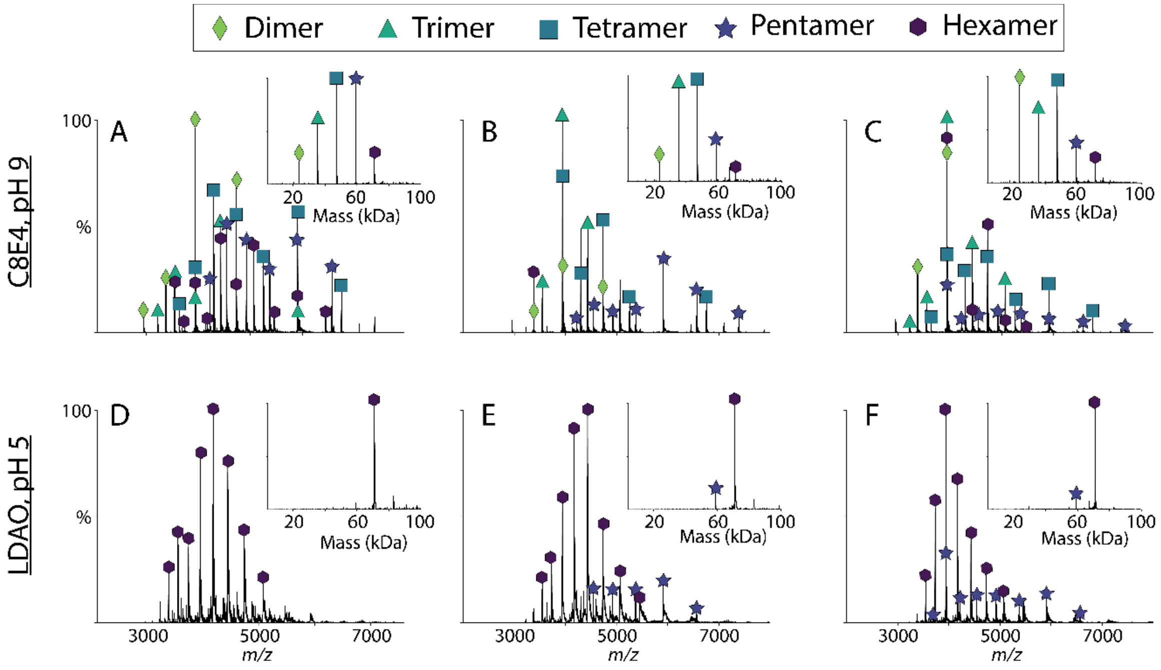

**Fig. S2:** Native mass spectra and deconvolved mass spectra (inset) of WT full-length AM2 (at 50 µM monomer) in C8E4 detergent at pH 9 (A–C) and in LDAO at pH 5 (D–F) shown for three separate replicates.

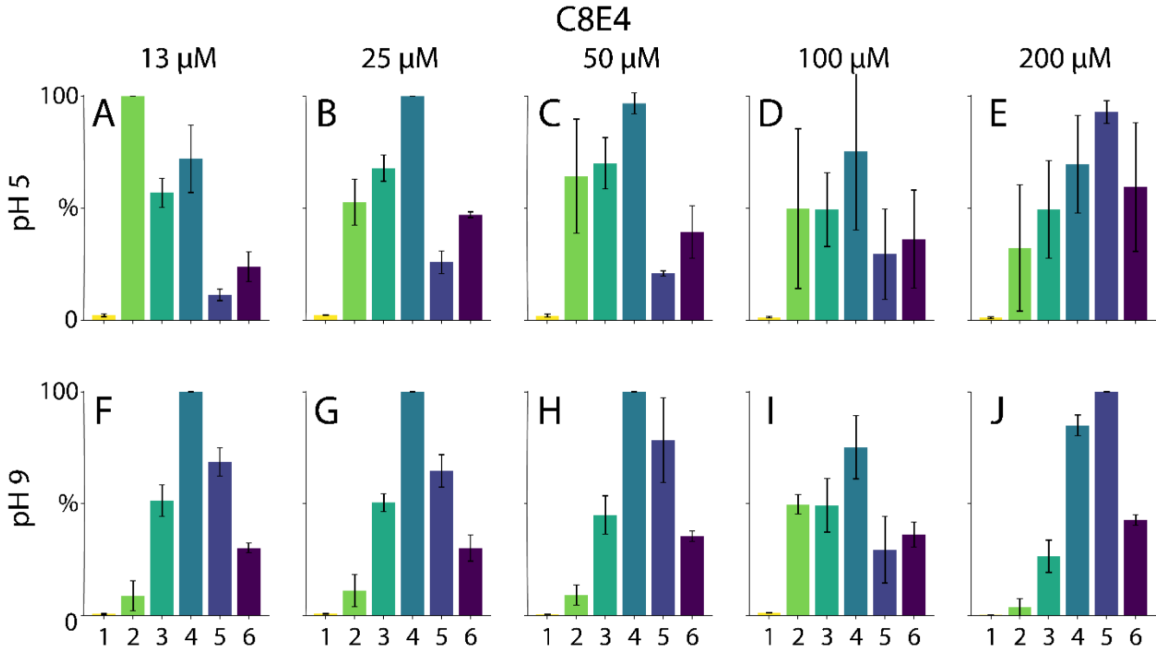

**Fig. S3:** The average relative peak areas measured by native MS of different oligomeric states of WT full-length AM2 solubilized in C8E4 at AM2 monomer concentrations of (A, F) 13, (B, G) 25, (C, H) 50, (D, I) 100, and (E, J) 200 µM at pH 5 (A-E) and pH 9 (F-J).

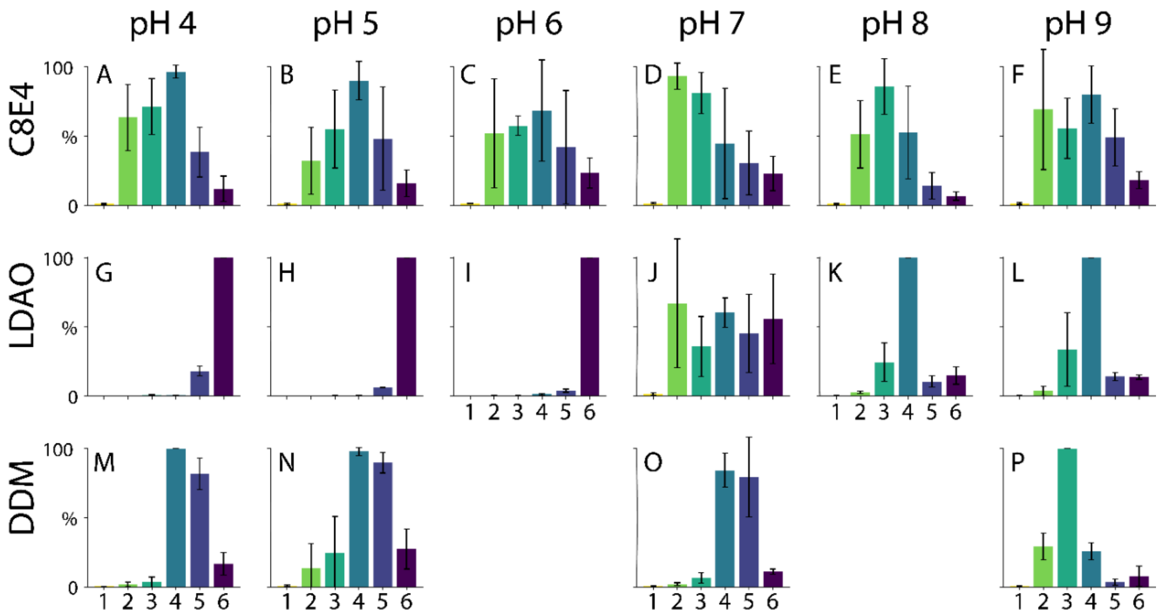

**Fig. S4**: The average relative peak areas measured by native MS of different oligomeric states of WT full-length AM2 (at 50 μM monomer) at pH 4 (A, G, M), pH 5 (B, H, N), pH 6 (C, I), pH 7 (D, J, O), pH 8 (E, K), and pH 9 (F, L, P) while solubilized in C8E4 (A–F), LDAO (G–L), and DDM (M-P). AM2 was not stable in DDM at pH 6 and 8, so no data is shown. Representative native mass spectra of AM2 in LDAO are shown in Figure 3.

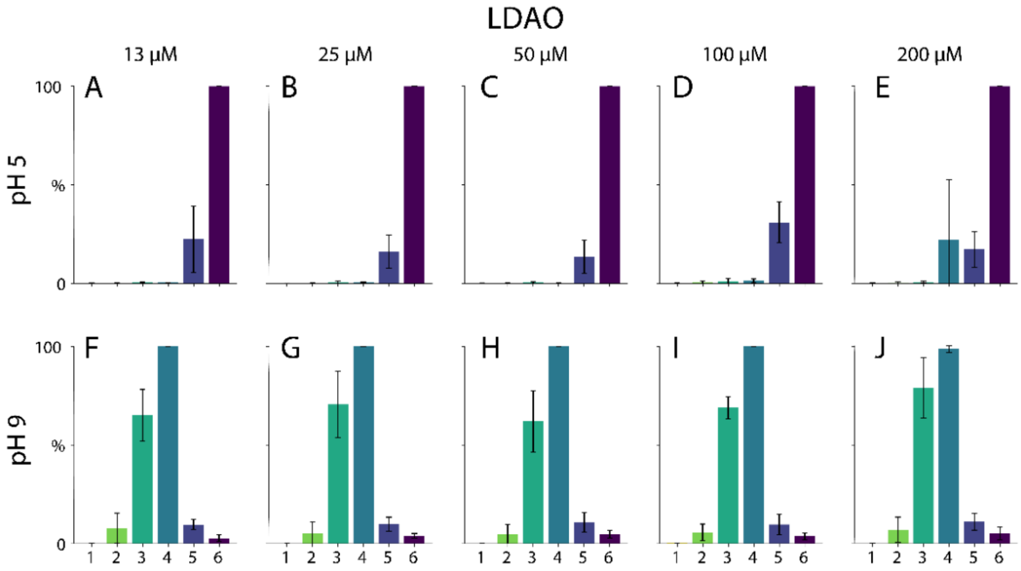

**Fig. S5:** The average relative peak areas measured by native MS of different oligomeric states of WT full-length AM2 solubilized in LDAO at an AM2 monomer concentrations of (A, F) 13, (B, G) 25, (C, H) 50, (D, I) 100, and (E, J) 200 μM at pH 5 (A-E) and pH 9 (F-J).

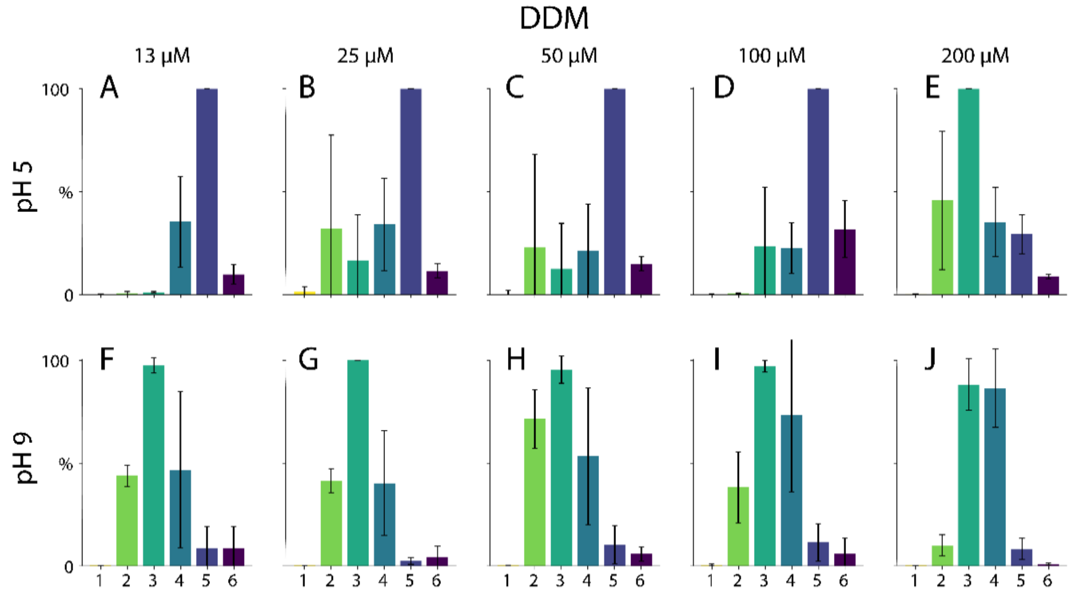

**Fig. S6:** The average relative peak areas measured by native MS of different oligomeric states of WT full-length AM2 solubilized in DDM at an AM2 monomer concentrations of (A, F) 13, (B, G) 25, (C, H) 50, (D, I) 100, and (E, J) 200 μM at pH 5 (A-E) and pH 9 (F-J).

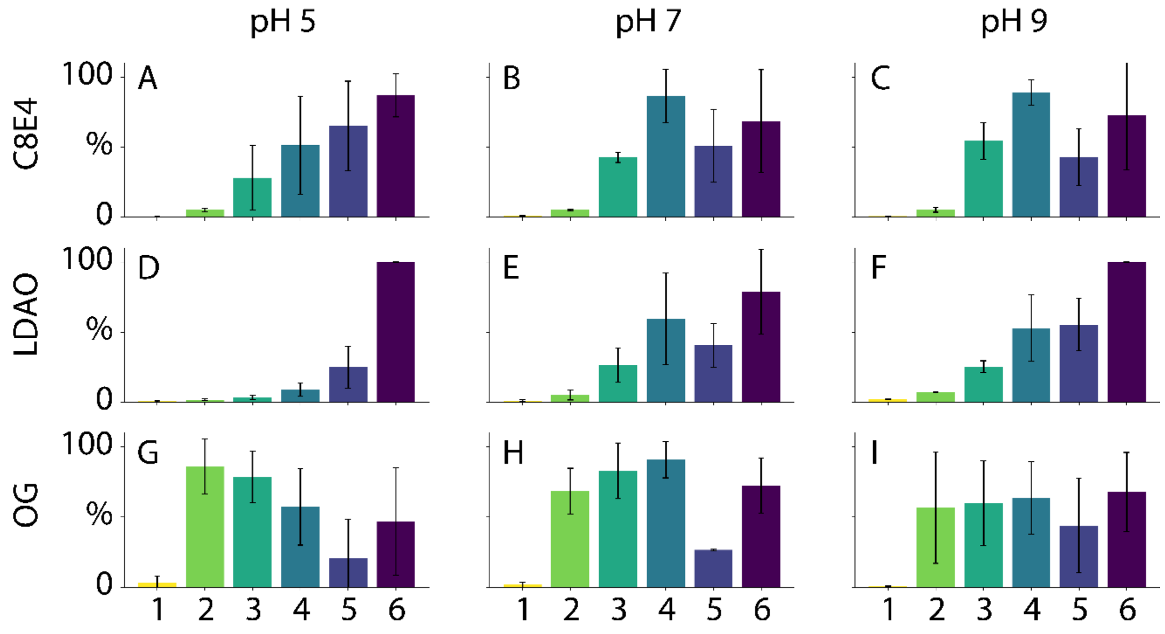

**Fig. S7:** The average relative peak areas of the different oligomeric states of the TM domain of WT AM2 (at 50 μM per monomer) at pH 5 (A, D, G), pH 7 (B, E, H), and pH 9 (C, F, I) while in C8E4 (A–C), LDAO (D–F), and OG detergents (G–I). Error bars indicate the standard deviation of measurements from triplicate samples.

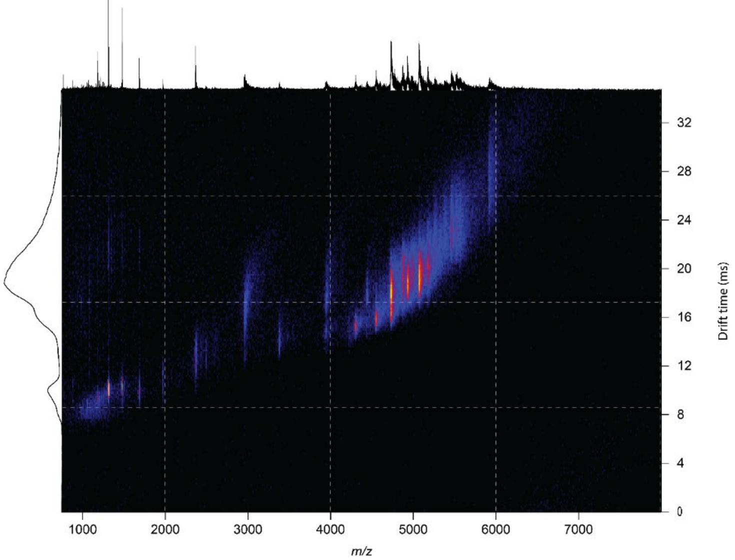

**Fig. S8:** Representative spectrum of IM-MS data for WT full-length AM2 in C8E4 pH 9 (at 50 μM monomer).

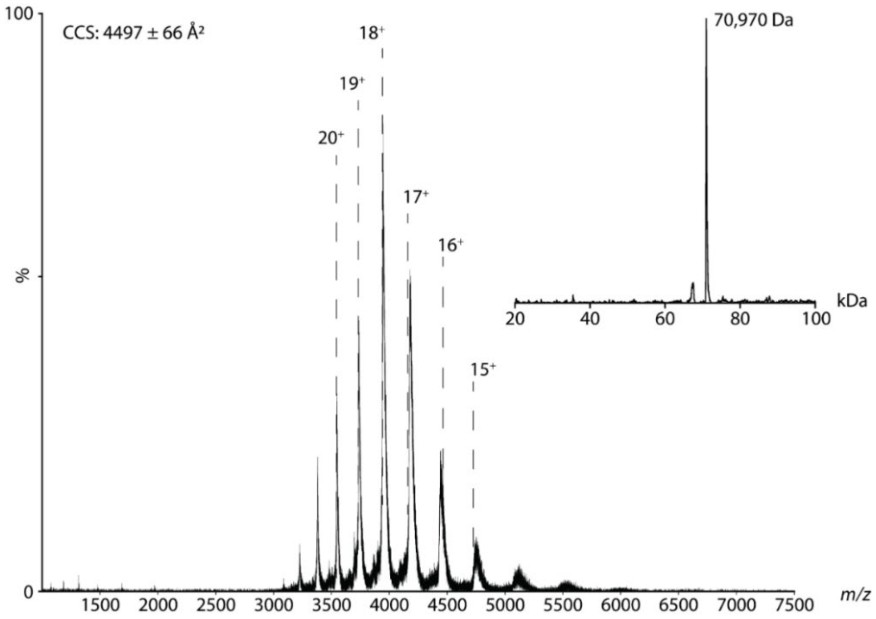

**Fig. S9:** Native mass spectrum measured with the Synapt XS Q-ToF mass spectrometer of WT full-length AM2 (at 50 μM monomer) solubilized in LDAO detergent at pH 5 with the deconvolved mass spectrum in the inset. Similar to results from an Orbitrap mass spectrometer (Figure 2D), a mostly monodisperse hexamer of AM2 is observed. The CCS value for the hexamer is 4497 ± 66 $Å^2$.

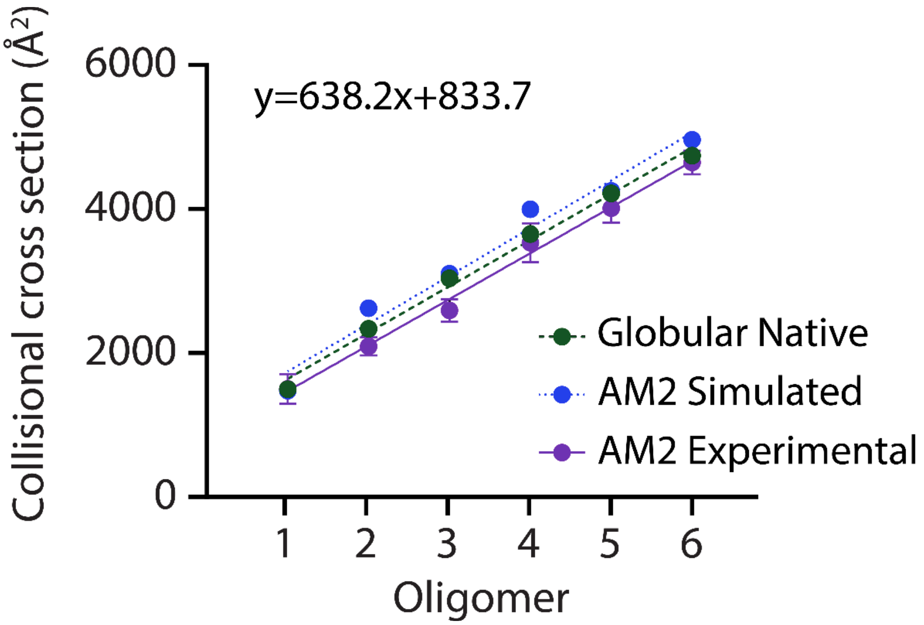

**Fig. S10***:* Experimental collisional cross sections (CCS) of AM2 oligomers in C8E4 at pH 9 (*purple*) compared to CCSs expected for native globular proteins (*green*) and to CCSs calculated for model structures (*blue*). A linear fit to the experimental data is annotated and shows that each monomer added to the oligomeric complex contributed around 638 $Å^2$ in CCS.

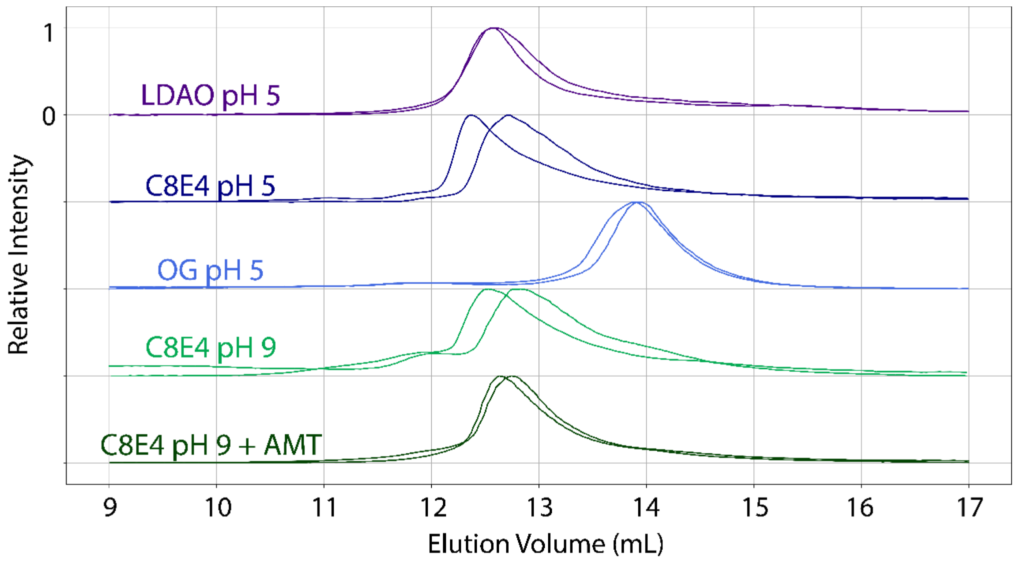

**Fig. S11:** The relative absorbances at 280 nm of AM2 in LDAO pH 5, C8E4 pH 5, OG pH 5, C8E4 pH 9, and C8E4 pH 9 with 300 μM amantadine during size exclusion chromatography. For comparison, standards were analyzed on the same column: thyroglobulin (eluted at 9.2 mL), catalase (9.6 mL), alcohol dehydrogenase (12.96 mL). carbonic anhydrase (16.39 mL), and ribonuclease A (17.59 mL). Duplicate injections are shown for each.

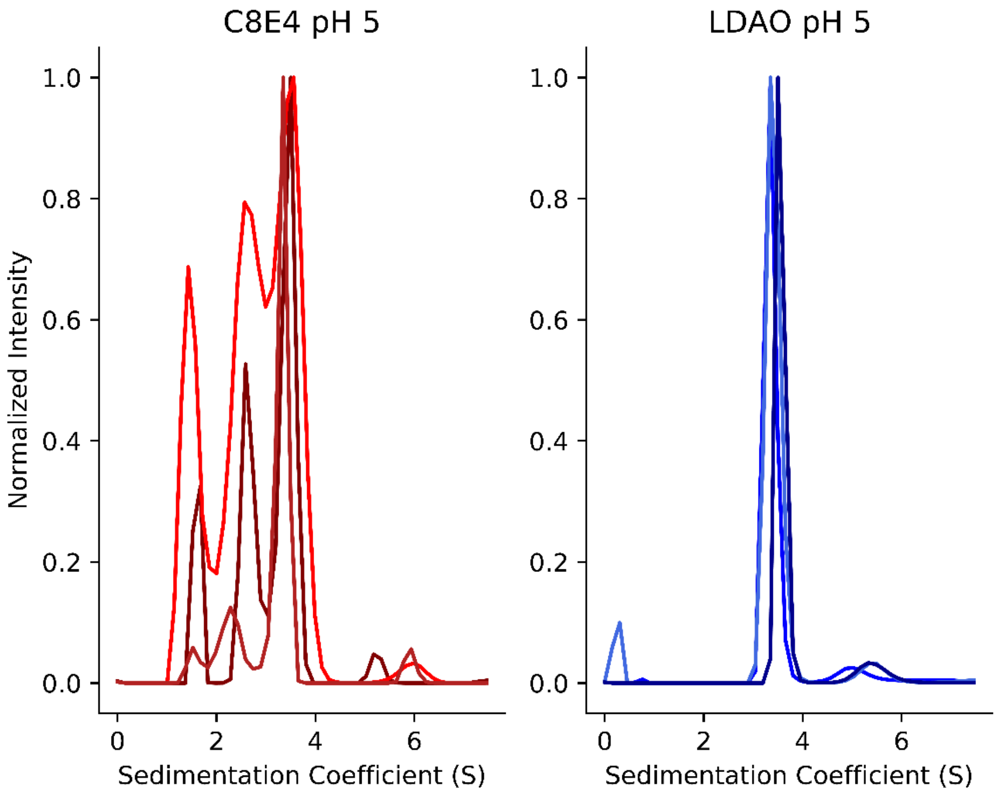

**Fig. S12:** The fit sedimentation coefficients (s20,w) from AUC of WT full-length AM2 solubilized in LDAO and C8E4 at pH 5 shown in triplicate.

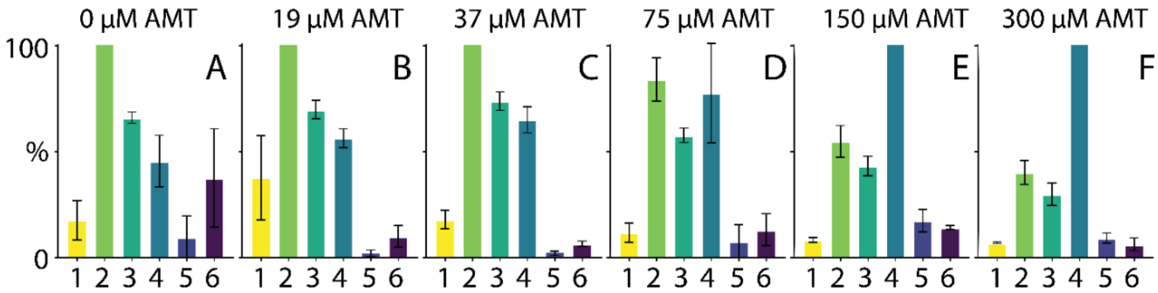

**Fig. S13:** The average relative peak areas of different oligomeric states of WT full-length AM2 in C8E4 (at 50 μM monomer) at pH 9 with increasing concentrations of amantadine added measured with the Synapt XS Q-ToF mass spectrometer. Increasing concentrations of drug drive formation of more monodisperse tetramer complexes.

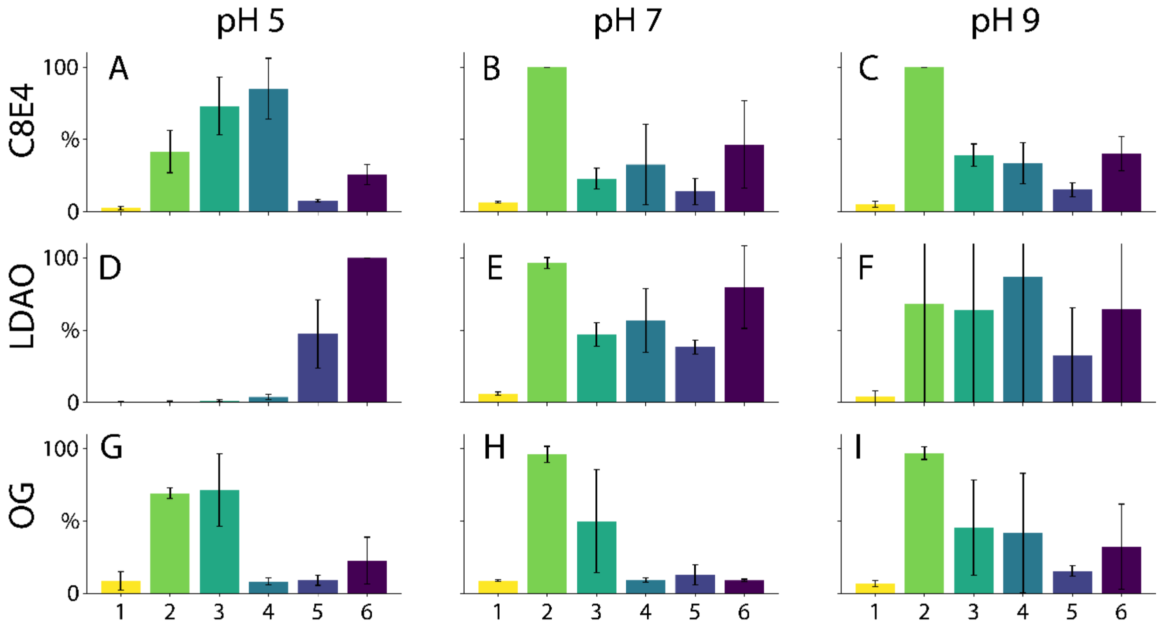

**Fig. S14:** The average relative peak areas measured by native MS of different oligomeric states of full-length AM2 S31N (at 50 μM per monomer) at pH 5 (A, D, G), pH 7 (B, E, H), and pH 9 (C, F, I) while solubilized in C8E4 (A–C), LDAO (D–F), and OG (G–I). Error bars indicate the standard deviation of measurements from triplicate samples

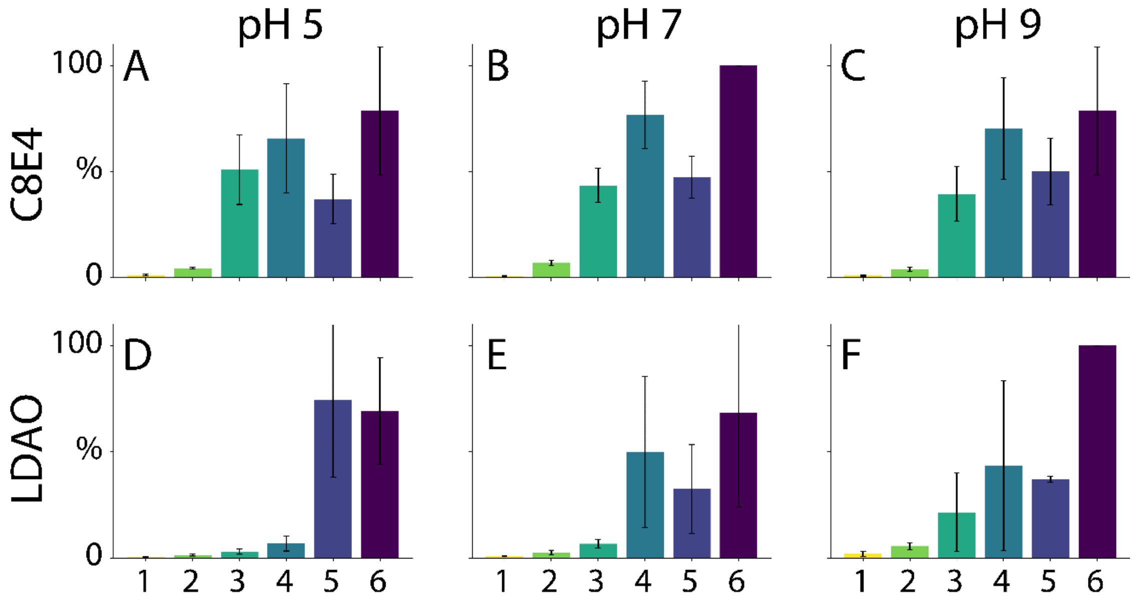

**Fig. S15:** The average relative peak areas of the different oligomeric states of the TM domain of AM2 S31N (at 50 μM monomer) at pH 5 (A, D), pH 7 (B, E), and pH 9 (C, F) while in C8E4 (A–C) and LDAO detergents (D–F). The error bars indicate the standard deviation of measurements from triplicate samples.

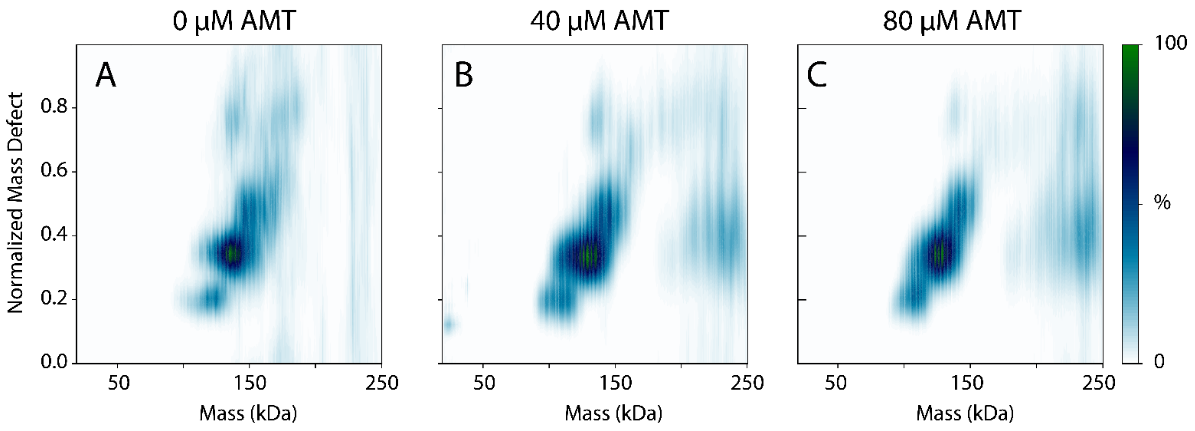

**Fig. S16:** Native MS intensities as a function of normalized mass defect versus mass for S31N full-length AM2 in DPPC nanodiscs with (A) 0 µM, (B) 40 µM, and (C) 80 µM amantadine.

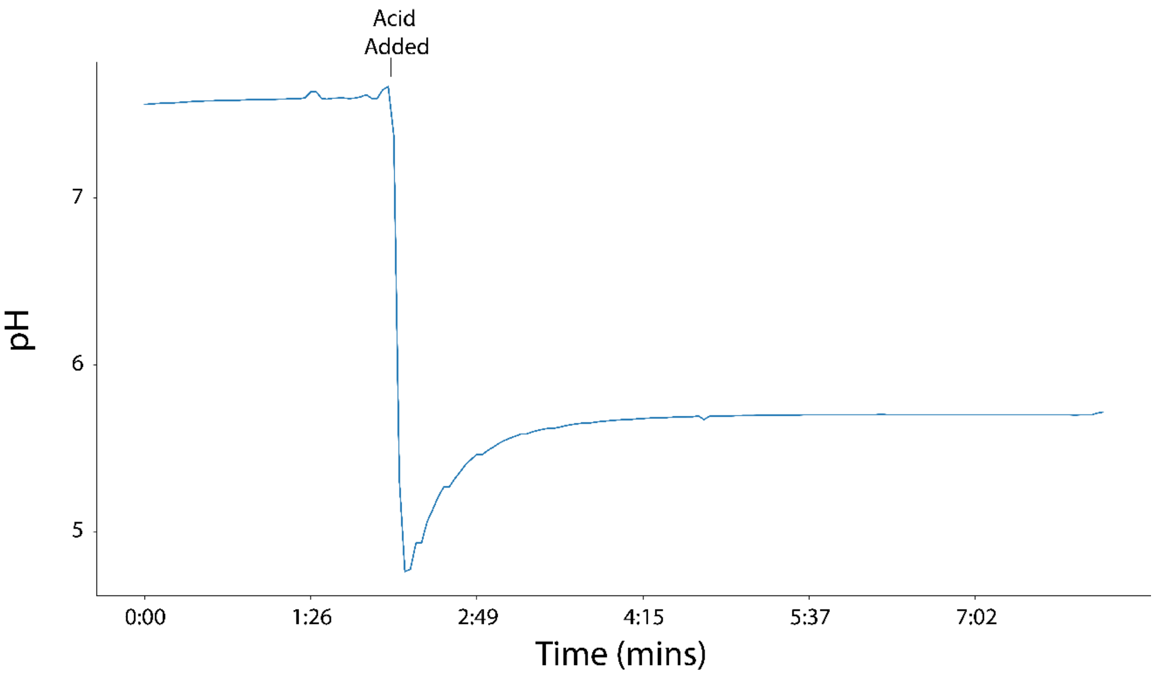

**Fig. S17:** Liposomal proton flux assay of WT full-length AM2 embedded within POPC liposomes. The assay was performed by adding acid to the external solution. The pH was measured as protons were transported by full-length AM2 to the interior of the liposome, which increased the pH of the external solution.

**Supplemental Tables**

**Table S1:** The theoretical masses, mean measured masses, and standard deviation of the mass measurement of AM2 in C8E4, LDAO, and OG detergents at pH 5 and 9.

| Theoretical Mass (Da) | Mean Measured Mass (Da) | Standard Deviation (Da) |
|---|---|---|
| 11810 | 11821.0 | 19.8 |
| 23620 | 23616.7 | 0.5 |
| 35430 | 35447.9 | 20.9 |
| 47240 | 47280.8 | 12.8 |
| 59050 | 59090.6 | 13.2 |
| 70860 | 70940.2 | 22.1 |

**Table S2:** Mass defect values for nanodiscs with different stoichiometries of AM2 that contain 2 × 22044 Da MSP belts with DMPC, DMPG, or DPPC lipids.

| AM2 Stoichiometry | DMPC | DMPG | DPPC |
|---|---|---|---|
| **0** | 0.03 | 0.09 | 0.09 |
| **1** | 0.44 | 0.81 | 0.16 |
| **2** | 0.86 | 0.51 | 0.25 |
| **3** | 0.28 | 0.22 | 0.34 |
| **4** | 0.70 | 0.92 | 0.43 |
| **5** | 0.12 | 0.63 | 0.51 |
| **6** | 0.54 | 0.34 | 0.60 |

**Table S3:** Mass defect shifts for binding different stoichiometries of amantadine in nanodiscs made of DPPC lipids. Contributions from AM2 and MSP are not included, so measured mass defect values will correspond to the values from Table S1 plus the shift indicated here.

| Amantadine Stoichiometry | 1 | 2 | 3 | 4 |
|---|---|---|---|---|
| **Mass Defect Shift** | 0.21 | 0.41 | 0.62 | 0.82 |
