## Supplementary material for "Influenza A M2 Channel Oligomerization is Sensitive to its Chemical Environment": Response to Reviewers

**Professor Michael T. Marty**
Department of Chemistry & Biochemistry
University of Arizona
Tucson, Arizona 85721 USA

THE UNIVERSITY OF ARIZONA TUCSON ARIZONA

Dear James,

We are pleased to submit the revisions for "Influenza A M2 Channel Oligomerization is Sensitive to its Chemical Environment" by Julia Townsend, Henry Sanders, Amber Rolland, Chad Park, Nancy Horton, James Prell, Jun Wang, and Michael Marty.

We thank the reviewers for their thoughtful comments and thorough review, which have greatly improved the manuscript. We have made a number of changes in response to reviewer suggestions, with detailed explanations below. These changes included a large number of new experiments, including orthogonal AUC data to confirm the oligomeric state trends observed with native MS, liposome assays to demonstrate protein function, and roughly doubling the amount of native MS data by including new experiments on the TM peptide and the S31N mutant in detergents. Overall, these new experiments largely confirm our initial findings that AM2 oligomerization is sensitive to its chemical environment, but they have also added additional nuance to the findings that we discuss.

**Reviewer 1:**

*1.1 How were the equilibration time and temperature of the samples for native-state MS analysis chosen? These two parameters (among others) can have significant effects on the population distribution of oligomers observed.*

All samples were allowed to equilibrate at room temperature for a few minutes. No differences in the oligomeric state were observed based on differences in equilibration time, even up to several days or incubating at different temperatures. We added a note describing this in the text.

*1.2 Page 5, first paragraph. "The precise oligomeric state distribution varied substantially between replicate measurements, indicating variable and relatively nonspecific oligomerization.".*

a. *Could the authors provide some context/examples on this variation between replicates? For most figures, a representative spectra or an average with error bars (with no individual data points noted) are presented.*

We have added a figure to show the differences between replicates for several conditions.

b. *Could the authors comment what implications the observed replicate variability would have on their interpretations of AM2 polydispersity?*

We thank the review for suggesting this and have added a discussion in the text discussing the implications of the variability. The variability is only seen in highly polydisperse conditions. More monodisperse samples are highly reproducible with very little variability.

*c. Could the authors explain why they conclude that the oligomerization is driven by relatively non-specific interactions? Prior structures of AM2, at least of the tetramer, show a symmetric oligomer with specific contacts being made at the interface between the monomers to form a conducting pore. Would the authors expect the interactions in the non-tetrameric states to be similar to or different from those observed in the tetramer?*

It could be that specific interactions (for example salt bridges between specific residues on adjacent monomers) still give rise to non-specific oligomers, where the same contact could be arranged at slightly wider angles to accommodate additional monomers in the complex. We could speculate that the contacts between the oligomers are similar, but without addition structural information, our data is not able to comment on this directly. We were meaning "non-specific" in relation to their oligomeric states, but we acknowledge that this invites readers to see these as driven by "non-specific" interactions, which we cannot determine in either direction from our data. We have changed some of the wording to "non-selective" avoid potential confusion.

*1.3 The fact that the replicate measurements showed significant variation suggests that these results may be due to technical complications rather than truly reflecting distinct complex formation. Did the authors consider using a positive control - perhaps something else known to form a tetrameric complex of similar molecular weight for comparison? This would help build confidence that utilizing native MS for this application can provide reliable data.*

There was only significant variation in samples that were very polydisperse. Conditions that appeared to form specific complexes (ex. LDAO pH 5) had very little variation between samples. We believe that the variation in the polydisperse samples captures the dynamic nature of these samples and sensitivity to subtle chances in sample conditions.

As positive controls, we have added a reference to our prior published work on streptavidin, which forms a tetrameric complex of very similar molecular weight. Here, only a tetramer is observed. Similarly, prior work on the same instrument with a small dimeric membrane protein, semiSWEET, with nearly the same monomer mass also showed clearly specific complexes, and we have referenced that.

*1.4 Were oligomers/aggregates larger than hexamers observed?*

Occasionally, in the most polydisperse samples (C8E4), some 7-mers and 8-mers were observed at much higher concentrations, but not at the working concentrations typically used (50 µM). For simplicity, we have not included these low abundance complexes in our discussion.

*1.5 In Figure 4, the distribution with 0 uM AMT of WT AM2 solubilized in C8E4 appears quite different than in Figure S1 and in the Figure S9 QToF data. Could the authors comment on the reproducibility of these distributions?*

The variation between Figure S1 and Figure 4, collected on the same instrument by different weeks, reflect the variability between samples as discussed above, which we expect is driven by the polydisperse nature of the sample that responds to subtle differences in the day-to-day sample preparation.

The differences with Figure S9 likely also reflect day-to-day variations in sample prep but could also reflect different biases in the instrumental transmission. For example, the Q-ToF appears more biased towards smaller ions, which would cause the lower oligomers to have higher relative signals. Because of these potential transmission biases, we have been careful to avoid any conclusions based on this information, as we mentioned in the text.

For more monodisperse samples (Figure S7 and S3), the results are highly consistent between days and between instruments, and these overall trends were highly consistent and reproducible between both instruments.

*1.6 Monomeric AM2 appears to be very low or non-existent in detergent, but is present in nanodiscs. Could the authors comment on how the detergent vs nanodisc environment could be responsible for the observed differences?*

We do not have a clear explanation for this interesting point. We agree that the detergent experiments generally show less monomer than the nanodiscs experiments, but there is some monomer present in a few detergent conditions (primarily OG and DPC) and no monomer present in DMPC nanodiscs. Interestingly, the new data from the transmembrane domain peptide show much less monomer in nanodiscs, so the monomer seems mostly unique to the full-length protein.

*1.7 Did the authors investigate the dependence on the AM2 to nanodisc ratio on the oligomeric distribution of AM2?*

Yes, as part of our initial experiments, we tested several different ratios. We have added this sentence to the manuscript to describe the results: "Lower ratios of incorporation showed less AM2 incorporated and higher ratios showed complex spectra that were difficult to resolve and interpret."

*1.8 The authors suggest that the S31N mutant is unable to bind amantadine because it is locked in a predominantly non-binding pentameric state (based on Figure 4 data). However, in nanodiscs, the S31N mutant forms monomers/dimers/trimers but no larger oligomers. Could the authors comment on this observed difference in their data, and how the authors' proposed mechanism of resistance relates to previous studies on the mechanism of the S31N mutant?*

To address this question, we performed native MS on the S31N mutant in a variety of detergent conditions. The more extensive data set showed more similarities between the WT and S31N mutant, so we have revised our conclusions.

*1.9 Page 9: "Importantly, AM2 S31N nanodiscs did not show any mass defect shifts upon addition of amantadine, confirming specificity of drug binding." Could the authors include this data, potentially in the supplementary file?*

We have included this data as a supplemental figure.

**Reviewer 2:**

*2.1 According to the materials and methods section, the protein was analyzed at a concentration of 50 μM (of the monomer?), which is quite high. Understandably, if a tetramer is expected, then higher amounts of the monomer are needed. However, since the protein appears in a range of assembly states, non-specific oligomerization should be ruled out.*

In many conditions tested, protein concentration had little influence on the oligomerization. We refer the review to Figures S3, S4, and S5, which tested different concentrations. In some cases, we do think the binding is relatively weak/non-specific, and we have added a note indicating this in the manuscript. However, even some polydisperse conditions were not majorly affected by protein concentration.

*2.2 In the few cases in which dilution experiments were performed the extent of dilution is not indicated, i.e. what are the starting and end concentrations.*

For all dilution experiments, the end protein concentrations were labeled on the figure and in the caption. As noted in the materials and methods, the protein was highly concentrated (580 μM) upon purification. Each Bio-Spin 6 column yielded slightly different concentrations after buffer exchange but were generally in the 300–400 μM range. Samples were then diluted to the final concentration listed in the figure.

*2.3 The data in Figs. 4, S1-S6 and S9 is processed, can the authors provide representative raw spectra, so the quality of data can be estimated.*

This would involve making over 75 new figures. We included representative data in Figures 1, 2, 3, and S7 and have added additional representative data in response to review comments above. We defer to the editor, but our suggestion is that interested readers review the raw data, which has all been posted publicly in Figshare in simple text file format.

*2.4 The discussion section should be extended, with emphasis on the biological relevance of the results. Like what is the composition of the natural host membrane? How can polydispersity in assembly states benefit the influenza virus? and their similarity to the membranes tested. Does any of the tested conditions mimic the natural environment of the host membranes? Can any conclusions be drawn as to the endogenous assembly state of AM2 in the host cells? In a structural and chemical point of view what is the mechanism in which pH or lipid content affect assembly?*

One of the potential benefits we discussed was the ability to expand to wider pores when triggered by acidic conditions in the endosome, participating with the viral uncoating process. We have also added a sentence to highlight differences in lipid compositions between different viral strains and different organelles as potential routes for biological regulation. However, we do not want to over-interpret our results beyond these points. We hope future studies can explore more the lipid compositions and AM2 oligomerization in different biological contexts.

*2.5 AM2 is post-translationally modified. Can the author comment on this aspect and how do they think it affects the assembly state distribution?*

No post-translational modifications were present on the sample used in these experiments. We are not aware that PTMs affect AM2 oligomerization (Hu, et al. Front. Microbiol. 2020), but this is an interesting topic for future research.

*2.6 In Figs 4, S1, S2 and S3 the concentration of Am2 is not indicated.*

We have added the concentration in the caption. As noted in the methods, all experiments were performed at 50 µM except where noted otherwise.

*2.7 The mass defect analysis should be explained.*

Additional explanation of mass defect analysis has been added to the supplemental methods.

*2.8 Raw data of the IM-MS results shown in Fig. S6 should be provided.*

Representative spectrum of IM-MS data has been added (Fig. SC).

*2.9 Theoretical and measured masses, including mass measurement errors should be added (also of drug binding). Perhaps in a table.*

A table with the masses has been added to the supplemental section. A table with the mass defect shift of drug binding was also added in the supplemental.

*2.10 What does the cartoon in figure 5 demonstrate?*

The cartoon was meant to show a schematic of directly ionizing AM2 in intact nanodiscs. To clarify this, we have added an explanation in the caption.

**Reviewer 3:**

*3.1 The author's note that "There are several dozen X-ray or NMR structures of the AM2 TM domain in a variety of membrane mimetics, all depicting monodisperse homotetramers" yet most of their conditions do not replicate this finding. Could the authors please comment in more detail on how their conditions differ from the previously reported structural studies which indicate AM2 is present as a homotetramer?*

*The authors mention that most studies used high concentrations of drug - are there other explanations as to why they observed high variability and complex instability where others did not? Do all the previous studies use drug to stabilize the complex? In cases where they did not use drug, what was different?*

We were also puzzled why our results did not agree with prior structures and have tried to rationalize this in several ways. Most studies do use added drugs, even when the drug might not be resolved in the structure. Also, most studies are done by NMR, where it is challenging to determine the oligomeric state distribution of homo-oligomeric complexes without advanced techniques that are often not employed. The high concentrations used in NMR may also push the complex towards tetramers. Furthermore, many studies have been done on only the transmembrane domain (TM) peptide. We repeated our experiments on the TM peptide and found that the TM can be more biased towards tetramers and dimers in membranes, although it also shows polydispersity in detergents.

Finally, structures are only published for conditions that are well-resolved, so our best guess is that some conditions tested for NMR or X-ray screening did show a polydisperse mixture but that these did not crystallize well or show well-resolved NMR spectra and thus were not pursued further by researchers. We aren't arguing that those structures were wrong, simply that there may be other states not sampled.

*3.2 In figure 2 and S1, please provide intensity values associated with each condition. Larger complexes are harder to ionize and more likely to inadvertently dissociate in the gas phase. It is impossible to understand how well AM2 ionized in each of these conditions when it is presented as a percent of total.*

The raw intensity values can vary substantially between replicate injections due to small differences in the disposable electrospray needles, where a different needle is used per injection. Thus, we are hesitant to include the intensity values, and we always normalize internally to the spectrum, which is conventional in native MS.

We agree with the reviewer that there could be biases in ionization and transmission based on their size. We have noted this in the text and been careful that our interpretation of the data takes this potential into account.

We are confident we are not seeing gas-phase dissociation. Gas-phase dissociation would almost certainly show asymmetric dissociation leading to highly charged monomers and lower charged oligomers. We do not see any evidence for these sorts of shifts, and we do not detect significant amounts of monomer. Similarly, gas-phase dissociation should lead to gas-phase unfolding that would be detected during ion mobility, which we do not observe. We have added a note to the text indicating this. We have also added additional orthogonal AUC measurements, which confirm the trends observed in native MS.

*3.3 Have the authors considered creating covalently bonded versions of dimer, trimer, and tetramer AM2 to use as standards to accurately quantify the amount of each complex in each condition?*

Creating covalently-bound complexes for calibration is an intriguing idea, but the amount of development and validation such an approach would take places it outside the scope of this manuscript.

*3.4 In figure S2, as protein concentration increases, a shift towards higher molecular weight complexes is observed. Is it possible this is due to protein aggregation and unlikely to be observed in physiological conditions?*

Figure S2 shows results for AM2 in C8E4, among the most polydisperse of all conditions tested. We believe that the protein is highly dynamic and does not form selective complexes under this condition, which explains the shift towards higher molecular weight complexes under higher concentrations. This trend is not observed under conditions where more selective complexes are observed. For example, Figure S4 shows that AM2 in LDAO does not form larger complexes at higher concentrations.

*3.5 The "orthogonal measurements confirm oligomeric sensitivity" section is confusing. What do the authors mean by oligomeric sensitivity? It is also unclear how the SEC data supports the authors' claims about the oligomeric state of AM2.*

We have changed the title and added text to clarify this section. The SEC data gives qualitative comparisons of the polydispersity of the sample through comparison of the overall peak shape under different conditions (more polydisperse samples had a broader peak shape, more monodisperse samples had a narrower peak shape). As noted above, we have also added AUC data as another orthogonal measurement technique.

*3.6 Could the authors please comment on why the select conditions were chosen for figure 2? Supplemental figure 1 is more informative and is worth including in the main figures. Similar question for the other figures where partial datasets are shown in the main text.*

Figure 2 highlights some of the most interesting conditions for AM2 that are discussed in detail in the discussion section. Figure S1 provides a more comprehensive look at all the conditions examined. We chose to include Figure 2 in the main text because it shows the raw data.

*3.7 Please clarify the concentration of AM2 used in Figures 1, 2, 3, 4 and S1 and S3.*

This information is provided in the supplemental methods. All measurements were at 50 µM for the monomer unless stated otherwise. We have added concentrations to figure captions for clarity.

*3.8 The authors offer two interpretations of their data in the discussion: 1) that it is very challenging to capture the pure tetramer 2) that the oligomeric states of AM2 are more complex than previously thought. The former is unlikely to have any physiological relevance while the latter could have important implications for development of novel therapeutics. A third interpretation could also be that the oligomeric profile observed is a byproduct of the native MS technique utilized. This manuscript would be much more impactful if this study included experiments to differentiate between these possibilities.*

We have used an orthogonal technique (analytical ultracentrifugation) of AM2 in different detergent conditions and found similar oligomeric trends, ruling out that our measurements are a byproduct of native MS.

**Reviewer 4**

*4.1 Why are the lipid nanodisc experiments only done at pH 7.4 and not other pH? In the detergent study, we clearly see a change in the oligomerization state brought on by a change in pH, and the authors speculate that the change in pH in the endosome could change the oligomerization state to higher order oligomers, so why is there no pH-dependent study of M2 in nanodiscs?*

This is an excellent suggestion that we tried but was unfortunately not successful. We are unable to assemble nanodiscs (even empty nanodiscs) at higher and lower pH. We also tried to exchange assembled nanodiscs into higher and lower pH, but these were also not stable.

*4.2 There have been several studies that look at the effects of a completely different set of detergents on the conformational landscape of the channel using solution NMR (Thomaston et al. JACS 2019) or different lipids using solid state NMR (Mandala et al. JMB, 2017): how does this study compare to these results? If the authors do the detergent study with solution state NMR, would they see evidence for polydispersity? Similarly, if the authors do these same native MS experiments using the detergents and/or lipids discussed in these two manuscripts, would they see polydispersity or do these conditions favor the exclusive formation of the homotetramer? The choices for lipids/detergents are orthogonal to what has been published in the literature, so a couple of experiments with the same sample conditions (i.e. lipid/detergent and pH) would be insightful as to whether the previous conditions just happen to favor the homotetramer.*

We have selected several conditions that AM2 has been previously studied in, including DMPC and DPPC bilayers, as well as DPC and OG micelles (PDB structures: 3BKD, 2MUV, 3LBW, 3C9J, 5C02, 2H95, 1NYJ, 2KQT). Our experiments with DPC and OG micelles at pH 7 match the NMR detergent experimental conditions most closely. The primary difference is that the NMR experiments were carried out at 2 mM peptide whereas ours were 50 uM. We do not see monodisperse tetramer under these conditions, perhaps due to the differences in concentration. However, our results from DPPC and DMPC are in good agreement with past results, especially with the TM peptide. With a few exceptions, our detergent results are generally inconsistent with past literature whereas the bilayer results are largely consistent. We have added a discussion of this to the text.

*4.3 In the amantadine-binding study of the WT and S31N in detergent micelles, the authors noted no major changes to the oligomeric state distribution for the mutant and conclude that the absence of a shift is indicative of lack of drug binding. They also suggest that the known drug resistance of the S31N variant arises because this mutant is locked into a novel pentameric state that is impervious to drug-binding. While this is an interesting hypothesis, their MS data does not prove that the drug is not binding. Moreover, they note that even in their WT samples, which show clear shifts, there is a lack of signal from the bound drug in their MS results, so how can the authors make the claim that S31N is not binding the drug? A similar comment can be made about the S31N nanodisc study, although the experimental evidence for drug-binding in the WT lends more support to this conclusion than the one made in the detergent study.*

We agree with the reviewer that our data cannot comment on drug binding in detergent, only on how the presence of the drug alters the oligomeric states. We have edited and added text to clarify this.

As the reviewer also notes, the binding is observed in the nanodisc experiments, so the presence of the positive control with the WT gives us confidence that the drug is not binding with the S31N mutant in those experiments.

*4.4 Can the authors rule out effects from the varying peptide:detergent ratios? Each of these samples was run at 2x CMC (seemingly standard in the native MS field) with a constant monomer concentration of 50 uM, which works out to very different peptide:detergent ratios. At the same peptide:detergent ratios, how do the distributions compare to each other?*

This is an interesting suggestion. The reviewer is correct that 2x CMC is standard in the native MS field. Higher concentration of detergent can lead to large signals from empty micelles that overwhelm the protein signals. However, all the pH and drug titration experiments were performed at the same protein:detergent ratio, so we can rule out any effects here. Also, the effect of protein:detergent ratio was indirectly tested with the protein titration experiments in Figure S2 and S4, which had the same detergent concentration but different protein concentrations. There was some effect in C8E4 at pH 5 but otherwise little to no effect.

*4.5 Since the higher order oligomers (i.e. hexamers) in LDAO seem stable, could they potentially crosslink these samples to get a low-resolution structure of the hexamer?*

This is an excellent suggestion, and we plan to follow up with structural characterization of the hexamer in future studies.

*4.6 Is there polydispersity evident in other detergents for S31N?*

Yes, similar trends of polydispersity were observed for AM2- S31N as well. We added new data showing this in the Supplemental Information.

*4.7 Previous studies (Ref #35 in this manuscript, for example) which look at the oligomerization of M2 using analytical ultracentrifugation used dodecylphosphocholine (DPC) micelles as the membrane mimetic. Using this particular detergent, the authors of the JMB publication showed that the monomer-tetramer equilibrium was cooperative in the presence and absence of the drug amantadine. Is there a reason why DPC was not used in this study? It would be interesting to see what distribution of states this technique captures in the detergent primarily used for the classical analytical ultracentrifugation experiments.*

We did use DPC for these studies, see Figure S1M-O. Our results in the absence of amantadine were not consistent with the previous studies, showing dimer and hexamer as prominent oligomers but overall, fairly polydisperse populations. The previous AUC studies were performed at much higher concentrations of DPC (which are unfortunately not possible with native MS) and slightly lower protein concentrations, preventing a more direct comparison.

However, we note that the monomer-tetramer-hexamer equilibrium is equivalent to the monomer-tetramer-octamer fit and that the fit is only slightly worse for a monomer-trimer-tetramer and monomer-trimer-hexamer model (Table 1, DOI: 10.1021/bi990720m). It could be that a more complex model not considered in this original work fits the data even better. At the least, this prior work lends equal support the presence of hexamers, which we often see in these results.

*4.8 Can the authors comment on why the drug-binding studies were only done in C8E4 detergent? How does the drug affect the distributions of the oligomers in other detergents? Would the larger hexamer observed in LDAO also bind the drug?*

As discussed in the text, drug binding studies were done in all conditions tested. The shift in oligomerization only occurs in C8E4 at pH 9. We believe that the dynamics and polydispersity of this conditions is likely related to this oligomeric remodeling. Unfortunately, the drug binding was not stable enough to be captured during native MS in detergent, so we cannot on drug binding, as detailed in our response above.

*4.9 The authors comment that the thickness and fluidity of the membrane is known to modulate M2 activity and suggest that these changes are due to a shift in the observed population of states in their discussion. Functional studies (i.e. liposomal proton flux assays) in the various lipids tested would be helpful to drive this point home. I would like to see how the activity of M2 changes in these lipids and how it relates to the distribution of states observed in the native MS.*

We have measured the bound drug to the WT in a lipid bilayer, suggesting this protein is properly folded/assembled. We have also performed liposomal flux assays to confirm the activity of this protein. We plan to do more extensive function studies in different lipid bilayer environments in future studies.

*4.10 There needs to be a thorough review of the literature - since the results here represent an effort to refute more than 100 publications on the M2 tetramer.*

We have added a more extended discussion of prior literature in response to this and other reviewer comments. Our goals are not to refute the prior literature. We also see tetramers under select conditions. Instead, we are presenting our data that shows a potentially more complex picture than previously appreciated. We were frankly surprised by the results and spent a great deal of time trying to reconcile what we were seeing with the past literature. This manuscript represents our best attempt to understand complex data that should have been simple.

*4.11 The authors commented on the bilayer thickness/saturation of DPPC as a potential reason for the tetrameric specificity of M2 in these conditions. Similar speculation into the chemical or physical properties of the detergents that give rise to the observed oligomeric distributions would be welcome.*

We looked into how the chemical and physical properties may relate to the oligomeric distributions and were unable to find any overall clear trends. For example, we compared the chain lengths of the detergents and found that detergents with similar chain lengths resulted in different patterns of oligomerization (i.e. C8E4 and OG). We also looked at detergent head groups and also found that similar head groups also resulted in different patterns of oligomerization (i.e. DDM and OG).

*4.12 Figure 2: Since the main take-home message from the figure is the deconvolved mass spectra, which clearly illustrate the polydispersity of the sample, it may help to flip the inset and the mass spectra or move the mass spectra to the supplemental. To someone who isn't in the field of native MS, the representative mass spectra are distracting and detract from conclusions illustrated in the deconvolved spectra.*

*Figure 3: A similar comment to the remarks made in Figure 2 can be made for this figure as well.*

We understand and appreciate this comment, but we felt the higher complexity of the raw data warranted a larger plot. The comparatively simple deconvolved masses are easier to interpret from the smaller plots.

*4.13 Figure 5: Is there a reason for the exclusion of S31N data? Since the drug-binding can be clearly seen in the corresponding WT samples, it would be better to swap out one of the WT-AMT figures (since they both are very similar) for one that shows the S31N with the drug even if no clear mass defect shift is seen. The two concentrations of AMT binding to WT is probably meant to show.*

We have added this data in a supporting figure.

We again thank the reviewers with their thoughtful comments to improve the manuscript.

Dr. Michael Marty
Department of Chemistry and Biochemistry
University of Arizona
Tucson, AZ 85721
